# Ovarian extracellular matrix mechanics regulate oocyte-follicle interactions during female reproductive aging

**DOI:** 10.64898/2026.03.31.715512

**Authors:** Xingyu Shen, Haiyang Wang, Gefei Cao, Yaelim Lee, Jin Zhu, Loo Chien Wang, Tianyun Zhao, Siok Ghee Ler, Radoslaw M. Sobota, Rong Li, Jennifer L. Young

**Affiliations:** Mechanobiology Institute (MBI), National University of Singapore, 5A Engineering Drive 1, 117411, Singapore; Institute of Modern Biology, Nanjing University, Nanjing 210008, China; Bioprocessing Technology Institute, 20 Biopolis Way, #02-01 Centros, 138668, Singapore; Institute of Molecular and Cell Biology, Agency for Science, Technology and Research (A*STAR), 61 Biopolis Drive, 138673, Singapore; Department of Biological Sciences, National University of Singapore, 16 Science Drive 4, 117558, Singapore; Department of Biomedical Engineering, College of Design and Engineering, National University of Singapore, 4 Engineering Drive 3, 117583, Singapore

**Keywords:** ovarian stroma, transzonal projections, TGF-β, infertility, stiffness

## Abstract

Female reproductive aging is associated with ovarian functional decline, leading to infertility. During aging, biochemical and biophysical changes in the ovarian extracellular matrix (ECM) occur, yet how these properties affect follicle growth and oocyte quality remains poorly understood. Here we describe spatiotemporal changes in the ovarian ECM with age using mass spectrometry, immunohistochemistry, and nanoindentation. While follicle stiffness remains unchanged, stromal matrix remodeling is associated with a ∼2.5-fold increase in stiffness. To understand how this increase in stromal stiffness affects age-related follicular dysfunction, isolated young follicles were cultured in soft and stiff hydrogels mimicking young and aged ovarian stromal stiffness, respectively. Higher stiffness leads to a decrease in granulosa cell (GC) proliferation, oocyte quality, and GC-oocyte interactions mediated via transzonal projections (TZPs). RNA-seq revealed TGF-β signaling as a major pathway affected by stiffness, and activation of TGF-β signaling through Mongersen treatment rescued TZP formation and oocyte quality in stiff matrix. These findings provide mechanistic insight into how changes in ECM mechanics contribute to ovarian aging functional decline and reveal potential therapeutic targets to counter fertility loss associated with tissue aging and fibrosis.

## Introduction

Female fertility declines progressively with age, resulting in an increased rate of spontaneous abortion and a reduced rate of conception (*1*, *2*). This age-related fertility decline is primarily attributed to a decrease in follicular quantity and quality (*3*). As the ovarian reserve pool is finite and unable to self-renew, developing strategies to preserve or restore follicular quality is a main approach to address age-associated fertility loss.

A multitude of factors at both the cell and tissue level contribute to follicular dysfunction during aging (*4*). In aged follicles, granulosa cells (GCs) exhibit reduced proliferation and increased apoptosis, whereas oocytes more frequently produce eggs with abnormal spindle morphology and chromosome number, as well as exhibit mitochondrial and metabolic dysfunction leading to reduced developmental potential (*3*, *5*). Exchange of metabolites and growth factors between oocytes and GCs are essential for proper follicle function; however, during aging, GC-oocyte communication is impaired, characterized by a reduction in transzonal projections (TZPs). This reduction in cell-cell interactions limits metabolic flux and causes a decline in oocyte-secreted factors that regulate cell proliferation, differentiation, and steroidogenesis (*6*, *7*). At the tissue level, the extracellular matrix (ECM) plays vital roles in ovarian function, including in follicle development and ovulation. Alterations to ECM composition, structure, and mechanics are widely acknowledged as a hallmark of ovarian aging (*8–10*), resulting in increased expression of fibrosis-associated factors and higher tissue stiffness (*11*, *12*).

Recent studies have demonstrated that antifibrotic drug treatment can ameliorate ovarian fibrosis and extend reproductive lifespan in aged mice, highlighting the therapeutic potential of ECM modification to enhance fertility (*13–15*). Indeed, culturing young follicles in hydrogels with tunable stiffness revealed that increased matrix stiffness impairs follicular growth and hormone secretion (*16*, *17*), establishing a causal link between mechanical properties and follicular development. However, the underlying mechanisms by which matrix stiffness results in follicular dysfunction remain unknown.

To understand compartment-specific regulation in the ovary, we spatially characterized ECM organization and mechanical alterations in fresh murine tissues during reproductive aging. We then applied *in vitro* hydrogel systems mimicking young and aged stromal stiffness for isolated follicle culture to understand how age-related mechanical cues direct follicle growth and oocyte quality. Our findings reveal that altered TZP formation, regulated by TGF-β signaling, is a key downstream process affected by stiff ECM. By perturbing TGF-β signaling with Mongersen, a SMAD7 antisense oligonucleotide, oocyte quality and function improved despite the stiff surrounding matrix. These findings have important implications in reproductive decline and provide a potential target for preserving follicle fitness in a stiff microenvironment that is characteristic of aged ovarian tissue.

## RESULTS

### Global and regional ECM remodeling during ovarian reproductive aging

We first carried out quantitative mass spectrometry (MS) to characterize global ECM alterations during ovarian aging in tissues from reproductively young (6-8 weeks) versus aged (14-16 months) mice. Because ECM proteins constitute only ∼1-1.5% of the total proteome (*18*), we applied an enrichment protocol that retains ECM proteins while reducing cellular protein components (*19*, *20*). MS identified > 4,000 proteins in total, and after annotation using the Matrisome Database (MatrisomeDB 2.0) (*21*, *22*), we identified 99 ECM proteins. In aged ovarian ECM, vitronectin (VTNC), collagen type VI (COL6A5/6), collagen type IV (COL4A4), and laminin (LAMA3) were all found to increase compared to young ECM (Fig. 1a). Downregulated ECM proteins with age include Fibrillin-2 (FBN2), collagen type II (COL2A1), and zona pellucida glycoproteins 1/2/3 (ZP1/2/3) (Fig. 1a).

**Fig. 1.**
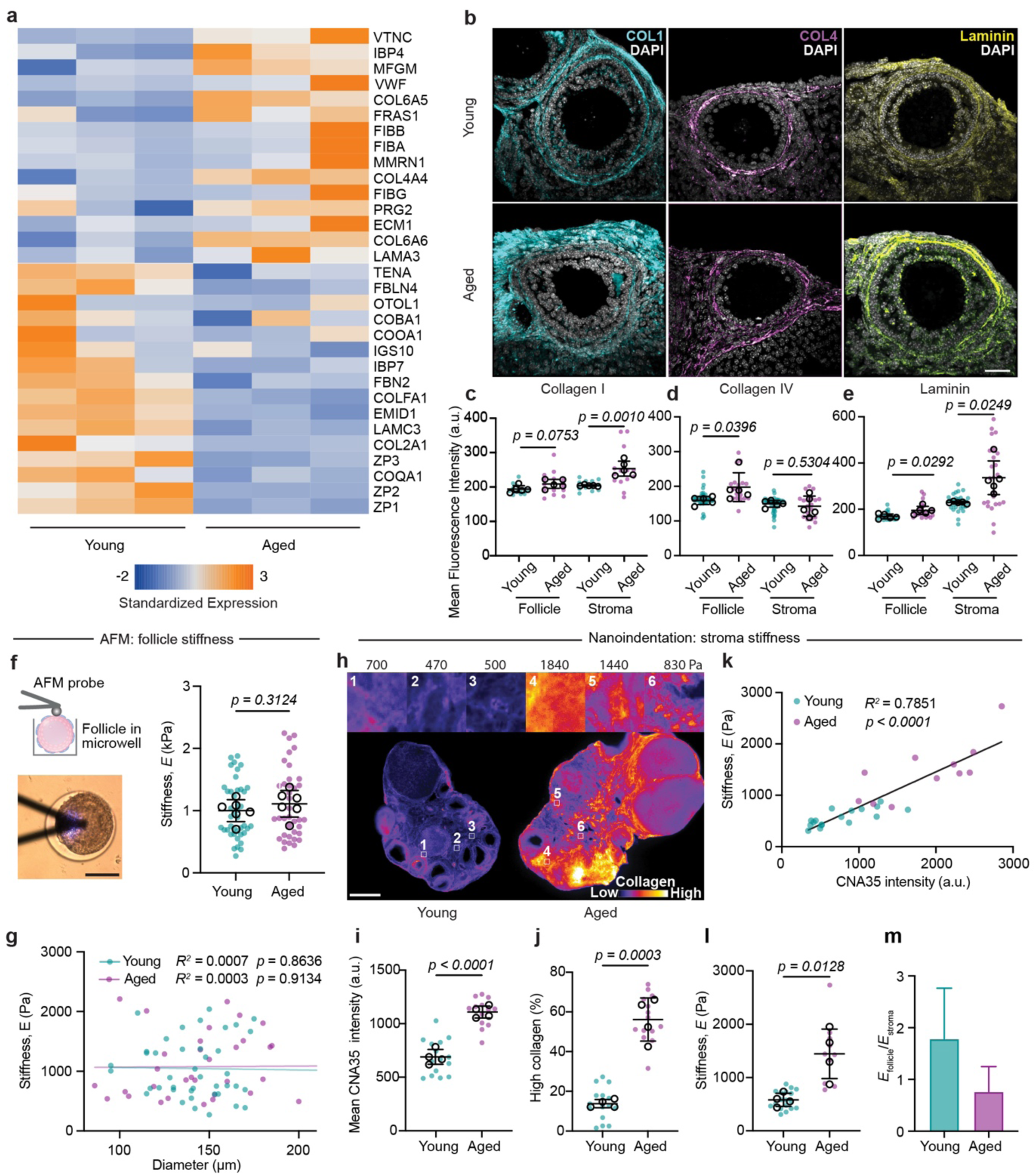
Age-related ECM alterations are spatially regulated and correlate with ovarian tissue mechanics. (a) Heatmap of the top 15 up- and downregulated core Matrisome proteins from MS of ECM in young and aged ovaries. (b) Representative immunostained images of collagen I, collagen IV, and laminin in young and aged ovarian tissue. (c-e) Mean fluorescence intensity for collagen I, collagen IV, and laminin in the stroma and follicle compartments. (f) Atomic force microscopy measurements of isolated young and aged follicles indented in a microwell, and (g) correlation between follicle diameter and stiffness. (h) Correlative collagen- stiffness measurements using CNA35 live probe and nanoindentation on fresh ovarian tissue slices. Representative images of CNA35-labeled ovarian slices from young (left) and aged (right) mice. Numbered white boxes indicate indentation areas with corresponding insets and Young’s moduli indicated (top row). Quantification of (i) mean CNA35 intensity and (j) percent collagen of high intensity (> 1000 px) in young and aged ovaries. (k) Correlation between tissue stiffness and CNA35 fluorescence intensity in young (teal) and aged (pink) ovaries. Each dot represents a single indentation. (l) Overall stiffness in young and aged ovaries, and (m) ratio of follicle to stromal stiffness with age. Sample sizes are: (a) N = 3 groups, with 4 - 5 mice per group; (b-e) 5 mice per group; follicle number for collagen I: n_young_ = 12 and n_aged_ = 14, collagen IV: n_young_ = 35 and n_aged_ = 18, laminin: n_young_ = 29 and n_aged_ = 24; (f-g) 6 mice per group, follicle number: n_young_ = 42 and n_aged_ = 37; (h-m) 4 mice per group, n_young_ = 16 and n_aged_ = 11. (c-g) teal and pink dots are individual follicles; black circles represent biological replicates. (i-l) teal and pink dots are ROIs; black circles represent biological replicates. Scale bars: 30 µm in (b), 100 µm in (f), 400 µm in (h). Statistical tests: data in c-f, i, j are shown as mean ± SD and analyzed using unpaired t-test. Scatter plot of g, k is analyzed with simple linear regression.

While bulk MS identified overall age-related changes in tissue composition, follicular and stromal ECM serve distinct structural and functional roles (*23*, *24*). Therefore, identifying where matrix remodeling occurs is essential to understand how these changes may influence follicle development. From our immunohistochemistry (IHC) images of collagen I, collagen IV, and laminin, which were identified as differentially regulated in our MS data, we characterized regional ECM abundance in follicular versus stromal compartments (Fig. 1b). With age, collagen I significantly increased in the stroma, collagen IV in the follicle, and laminin in both (Fig. 1c-e). As these proteins are known to affect ECM stiffness (*25*, *26*), their compartment-specific alterations with age likely contribute to spatially distinct mechanical properties within the ovary. Indeed, recent reports found age-related mechanical changes across ovarian compartments, specifically in the corpora lutea (*27*), and we next sought to understand the contribution of matrix to follicular versus stromal mechanics.

### Ovarian stromal but not follicle stiffness increases with age

To understand how follicular versus stromal ECM alterations correlate to age-specific mechanical properties, we carried out nanoindentation on both isolated follicles and fresh ovarian tissue sections. Atomic force microscopy (AFM) on isolated primary to secondary stage follicles from both young and aged mice was used to quantify follicle stiffness (Fig. S1a). We found no stiffness difference between young and aged follicles (Young’s modulus, *E* ∼1 kPa) nor a dependence on follicle size/stage (Fig. 1f, g). These findings are consistent with our IHC staining, where limited ECM remodeling was observed in the follicular regions (Fig. 1c-e).

For stromal mechanics, we used nanoindentation on fresh ovarian tissue sections that were live stained with CNA35-GFP, a collagen hybridizing peptide (*28*). This enabled us to correlate stromal stiffness with ECM organization by comparing indented regions with captured images of stained collagen (Fig. S1b). Overall, we observed an increase in collagen abundance in aged versus young ovaries (Fig. 1h-j), in line with age-related fibrosis (*10*). Correlation analysis revealed that stiffness is proportional to collagen content (Fig. 1k), with aged ovarian stroma being ∼2.5-fold stiffer than young stroma (Fig. 1l). Previous reports identified follicles as the mechanically dominant component of the ovary, which we confirm in young tissues (e.g., *E*_follicle_/*E*_stroma_ ∼1.7) (*29*) (Fig. 1m). With age, however, this relationship reverses (e.g., *E*_follicle_/*E*_stroma_ ∼0.8), consistent with ECM remodeling and suggesting that this shift in mechanical balance may contribute to follicular dysfunction (Fig. 1m).

### Matrix stiffness impairs follicle growth, GC proliferation, and oocyte quality

To understand how stromal mechanics impact follicle development, we produced alginate hydrogels mimicking young versus aged stromal stiffness for 3D *in vitro* culture of isolated secondary follicles from young mice (Fig. 2a, b). Consistent with previous work (*16*), follicles cultured in soft hydrogels over 7 days grew larger and had more proliferating Ki-67^+^ GCs than those cultured in stiff hydrogels (Fig. 2c-f). This reduction in proliferating GCs is a hallmark of aged ovaries, indicating the stiff hydrogel culture can recapitulate some aspects of aging dysfunction in young follicles (Fig. S2a, b). Next, oocyte quality was assessed and we observed a ∼3-fold reduction in maturation and a ∼2-fold increase in MII spindle abnormalities in stiff hydrogels (Fig. 2g-j). Taken together, these data show that stiffness alone can impair follicle growth, GC proliferation, and oocyte quality.

**Fig. 2.**
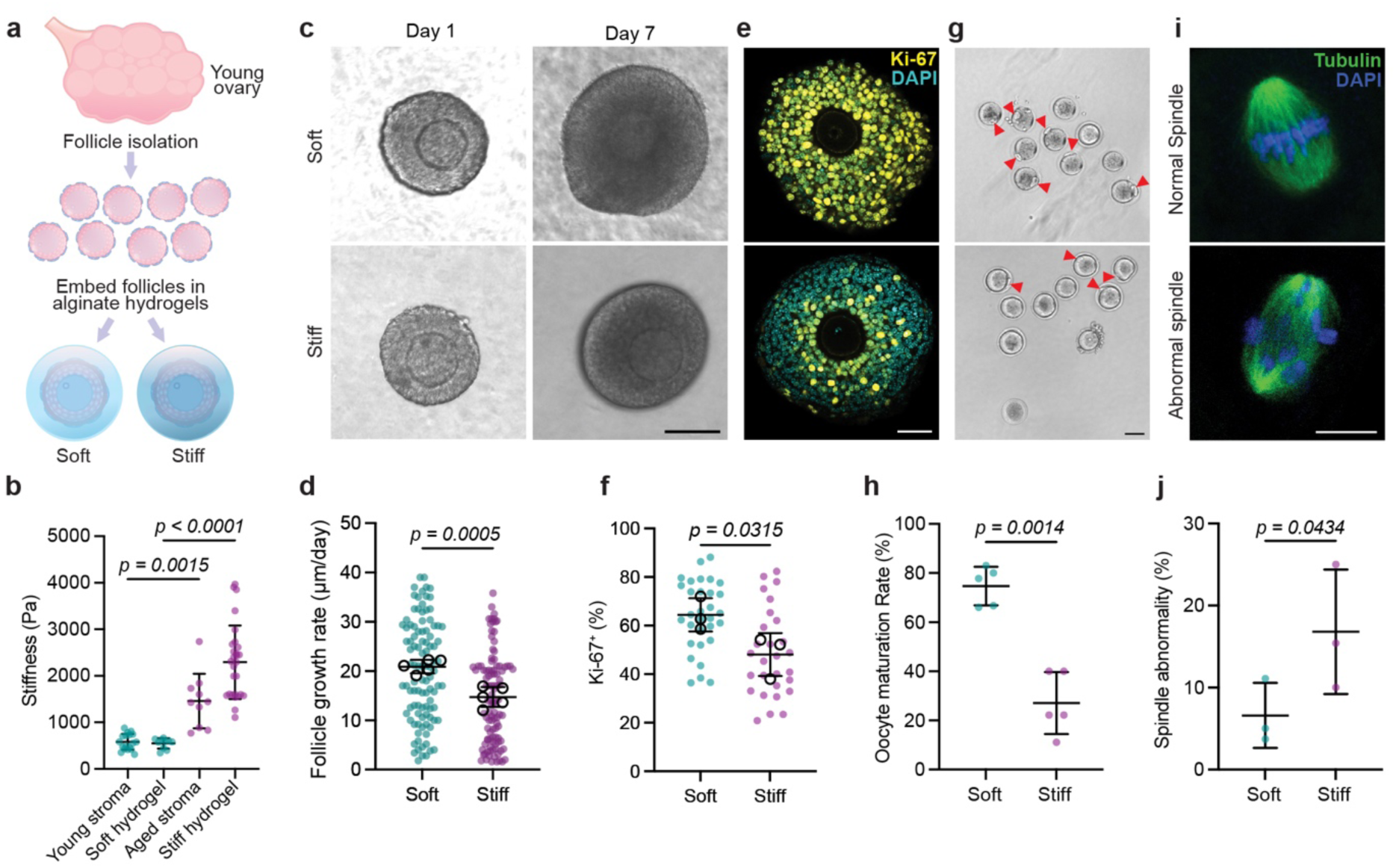
Matrix stiffness impairs follicle growth and oocyte quality. (a) Follicles were isolated from young ovaries and cultured in soft and stiff alginate hydrogels. (b) Hydrogel stiffness was tuned to that of young (‘Soft hydrogel’) and aged (‘Stiff hydrogel’) ovarian stroma based on our nanoindentation data. (c) Brightfield images of follicles grown in soft (top) and stiff (bottom) hydrogels over 7 days with corresponding (d) follicle growth rate (µm/day). (e) Ki-67 (yellow) and DAPI (cyan) immunostaining was carried out to assess proliferation in follicles cultured in soft and stiff hydrogels and (f) the percentage of Ki67-positive granulosa cells was quantified. (g) Oocyte maturation through polar body extrusion (red triangles) was assessed in brightfield post-*in vitro* ovulation and (h) quantified in soft and stiff hydrogels. (i, j) Spindle morphology was assessed as normal (top) or abnormal (bottom) using immunostaining of α-tubulin (green) and DAPI (blue) and quantified as % abnormality. (d,f) teal and pink dots are individual follicles; black circles represent biological replicates. (h, j) teal and pink dots are biological replicates. Data is shown as mean ± SD and analyzed using unpaired t-test. Sample size is: (b) n = 10 (soft) and 28 (stiff). (d) N = 5 replicates. n = 104 follicles in both soft and stiff hydrogels. (f) N = 3 replicates, n = 31 (soft) and 28 (stiff) follicles. (h) n = 5 independent experiments. (j) n = 3 independent experiments. Scale bars: 100 µm in (c), 70 µm in e and g, 20 µm in i.

### Stiffness alters both GC and oocyte gene expression

To understand how stiffness impairs follicle function, we performed RNA sequencing (RNA-seq) on separated GC and oocytes isolated from young follicles cultured in soft or stiff hydrogels (Fig. S3a). Differentially expressed gene (DEG) analysis of GCs identified a total of 1,160 DEGs, with 872 upregulated and 288 downregulated, in the stiff versus soft hydrogels (Fig. S3b, c). In stiff hydrogels, Gene Ontology (GO) analysis identified ‘ECM organization’, ‘response to mechanical stimulus’, ‘cell adhesion mediated by integrin’, ‘ERK1/2 cascade’, and ‘cellular response to transforming growth factor beta (TGF-β) stimulus’ as upregulated biological processes, while downregulated cellular components include ‘actin-based cell projections’, ‘adherens junctions’, and ‘cell projection membranes’ (Fig. S3d, e). DEG analysis of isolated oocytes from follicle culture in stiff versus soft hydrogels showed that global transcriptional differences were not as pronounced as those observed in the GCs (Fig. S3f); however, Gene Set Enrichment Analysis (GSEA) identified metabolic processes, including ‘mitochondrial gene expression’ and ‘NADH dehydrogenase complex assembly’, as downregulated with stiffness (Fig. S3g).

High matrix stiffness has been associated with the upregulation of specific mechanosensitive pathways, including the Hippo pathway, which functions to transduce mechanical signals to biochemical responses (*30*). Thus, we examined our GC RNA-seq data for *Yap1* and downstream Hippo pathway genes but found no differential regulation related to hydrogel stiffness (Fig. S4a). Consistently, immunostaining of YAP in follicles cultured in the soft and stiff hydrogels showed no significant difference in YAP nuclear translocation in the GCs (Fig. S4b, c). Collectively, these findings demonstrate that stiffness reprograms GCs and oocytes independently of canonical YAP signaling, suggesting the involvement of alternative mechanotransduction pathways contributing to follicular dysfunction.

### Matrix stiffness reduces TZP density between GCs and the oocyte

Further analysis of the GC RNA-seq data using GO terms ‘adherens junctions’ and ‘actin-based cell projections’ found that *Cdh2* (encodes N-cadherin), *Fmn2* (encodes Formin 2), and *Myo7a* (encodes myosin-VIIA) were significantly downregulated in stiff hydrogels (Fig. 3a-e). This led us to hypothesize that ECM stiffness affects TZPs, which are the cytoskeleton-based extensions from GCs to the oocyte that form both adherens and gap junctions between the two cell types. For follicles cultured in stiff hydrogels, we found a ∼28% reduction in TZPs per oocyte versus in soft hydrogels (Fig. 3f, g), in line with recent reports in other systems (*31*). This reduction in TZPs is attributed to a reduction in TZP density (defined as number of TZPs per 10 µm) rather than to oocyte diameter or the number of oocyte-adjacent GCs (Fig. 3h-j). Next, to determine whether these *in vitro* changes resemble those observed *in vivo* during aging, we quantified TZPs in directly isolated follicles from reproductively young and aged mice. Aged follicles had ∼50% fewer TZPs per oocyte, while oocyte diameter and oocyte-adjacent GC number also remained unchanged (Fig. 3k-o). Thus, this stiffness-induced TZP loss recapitulates the age-associated decline in TZP density.

**Figure 3.**
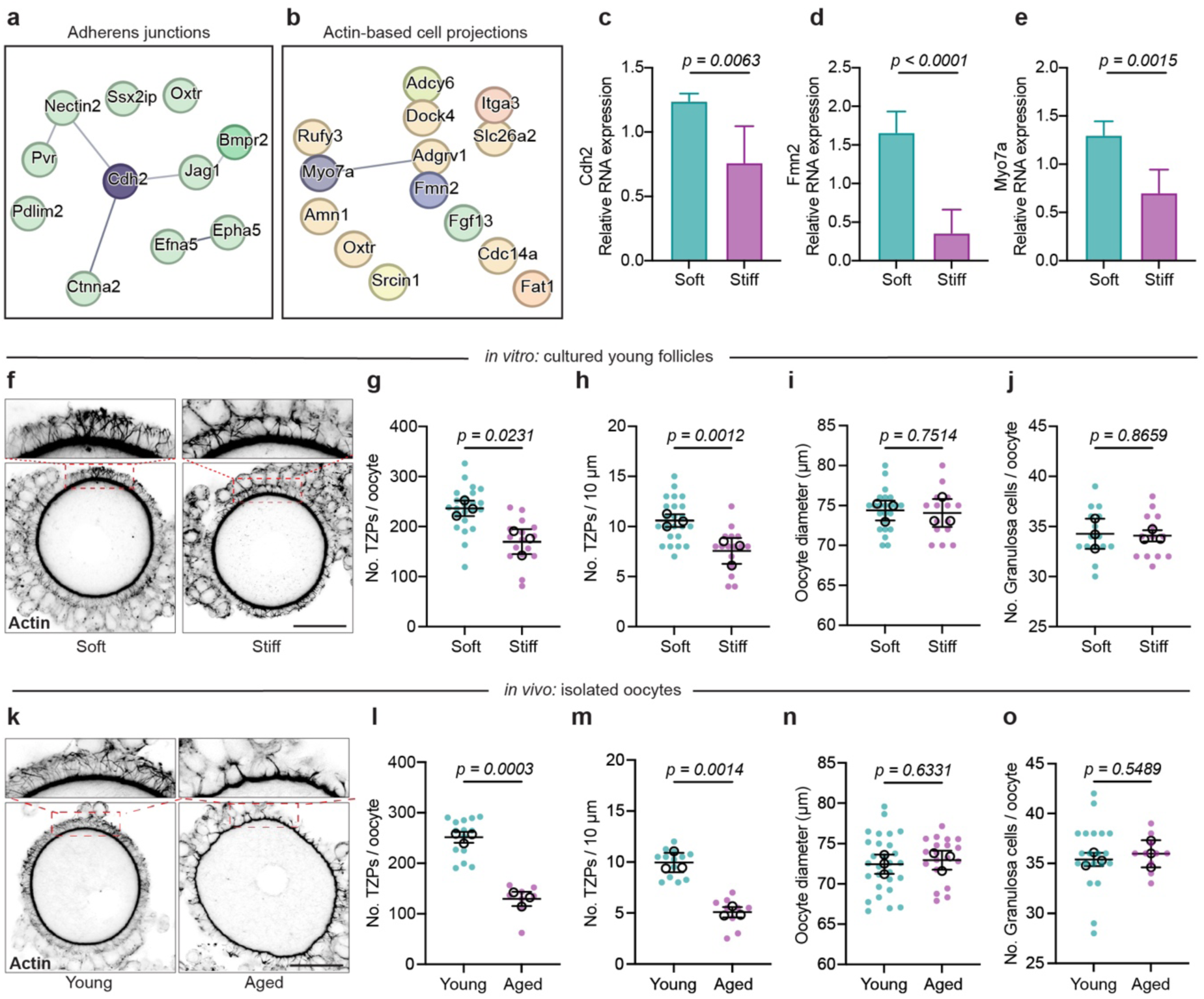
Matrix stiffness reduces TZP density, mimicking *in vivo* age-related phenotypes. (a-b) Protein-protein interaction (PPI) networks show down-regulated genes in GO terms ‘adherens junctions’ and ‘actin-based projections’ for follicles cultured in stiff vs. soft hydrogels. (c-e) Relative expression of genes Cdh2, Fmn2, and Myo7a in soft and stiff hydrogels. (f) Actin immunostaining shows TZPs in oocytes cultured in soft and stiff hydrogels with insets highlighting TZP disruption in stiff hydrogels. (g-j) Quantifications for total TZPs per oocyte (g), TZPs per 10 µm of oocyte perimeter (h), oocyte diameter (i), and granulosa cells per oocyte (j). To compare TZPs of *in vitro* cultured young follicles to *in vivo* values, (k) actin immunostaining was carried out on freshly isolated oocytes from young and aged mice, insets show TZP disruption with age. (l-o) Quantification was done as in (g-j). (g-j, l-o) teal and pink dots are individual oocytes; black circles represent biological replicates. Data is shown as mean ± SD and analyzed using unpaired t-test. Sample size is: n = 23 (soft) and 16 (stiff) oocytes in (g-i), n = 15 (soft) and 13 (stiff) in j, N = 3 independent experiments, n = 14 young and 9 aged oocytes in i-m, n = 28 young and 19 aged in n, n = 20 young and 10 aged in o. Scale bar in (f, k): 30 µm.

### Impaired TGF-β signaling underlies stiffness-induced TZP and follicle decline

GO analysis of GC RNA-seq data from follicles cultured in stiff versus soft hydrogels identified the ‘cellular response to TGF-β stimulus’ pathway as significantly upregulated (Fig. 4a). This pathway has previously been shown to regulate TZP formation and is mechanosensitive, although its mechanosensitive activation has not been demonstrated in ovaries (*32–35*). Thus, we investigated downstream regulators, specifically Smad7, which is a negative regulator of TGF-β signaling and competes with the Smad 2/3 and Smad1/5/8 complexes that translocate into the nucleus with co-mediator Smad4 to control gene transcription (*36*, *37*). Smad7 was found to be upregulated in stiff hydrogels (Fig. 4b), suggesting that the TGF-β signaling pathway may be inhibited by mechanics. Indeed, we observed a decrease in Smad2/3 nuclear translocation in GCs within the stiff hydrogel (Fig. 4c, d). When compared to *in vivo* tissues from reproductively young mice, Smad2/3 nuclear translocation was only observed in the GCs of large follicles, which was correlated to an increase in TZP density, consistent with activation of the TGF-β signaling pathway that has been reported to promote TZP formation (Fig. S5a-d).

**Figure 4.**
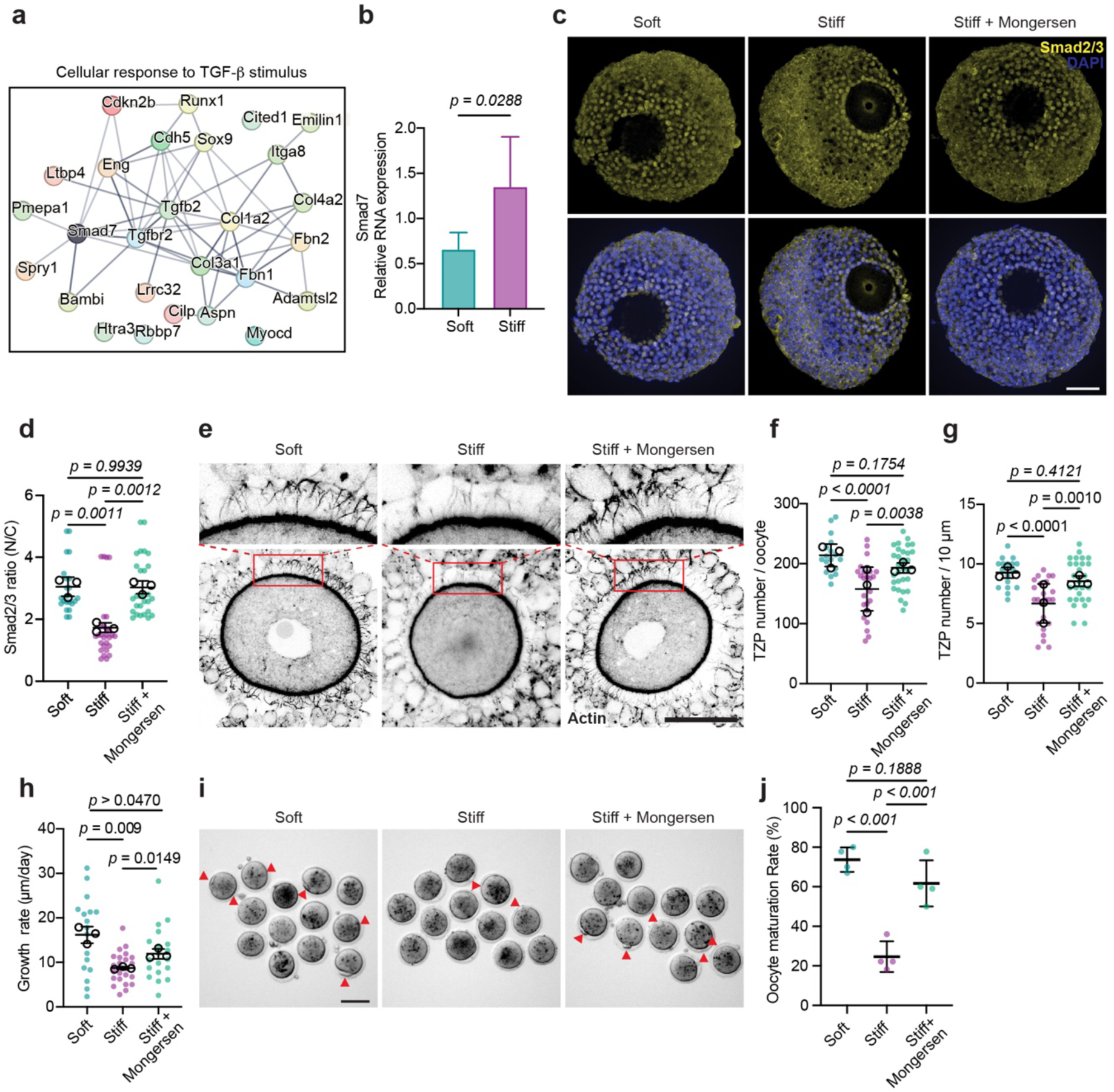
Activation of TGF-β signaling rescues follicle growth and restores TZPs in stiff hydrogels. (a) PPI analysis of DEGs from follicles cultured in stiff versus soft hydrogels identified the network ‘cellular response to TGF-β stimulus’. (b) Gene expression of Smad7 increased in aged follicles. (c) Immunostaining of Smad2/3 (yellow) and DAPI (blue) in follicles cultured in soft hydrogels, stiff hydrogels, and stiff hydrogels + Mongersen. (d) Nuclear translocation was quantified (N/C: nuclear to cytoplasmic ratio). (e) Actin immunostaining for TZPs in the soft, stiff, and stiff + Mongersen groups with insets with corresponding (f, g) quantification of TZP number and density. Functionality in these three conditions was assessed over 6 days of culture through (h) follicle growth rate (µm/day) and (i, j) oocyte maturation. (i) Brightfield images show polar body extrusion (red triangles) post- *in vitro* ovulation after culture in soft hydrogels, stiff hydrogels, or stiff hydrogels + Mongersen, with (j) quantification. (d, f-h) teal, light teal, and pink dots are individual follicles; black circles represent biological replicates. (j) teal, light teal, and pink dots represent biological replicates. Data is shown as mean ± SD and analyzed using One-way ANOVA. Sample size is: (d) n = 12, 14, and 14 in soft, stiff, and stiff + Mongersen groups, respectively. (f, g) n = 25, 24, and 30 in soft, stiff, and stiff + Mongersen groups, respectively. (h) n = 17, 23, 19 in soft, stiff, and stiff + Mongersen groups, respectively. N = 3 replicates. (i) N = 4 replicates. Scale bar: 50 µm in (c) and (e), 70 µm in (i).

To test if Smad7 knockdown could restore TZPs and follicle growth in stiff hydrogels, we treated our cultured follicles with Mongersen, a SMAD7 antisense oligonucleotide (*38*, *39*). Treatment increased Smad2/3 nuclear translocation in the stiff hydrogel, to the same level as in the soft hydrogels, suggesting TGF-β pathway activation (Figure 4c, d). Further, we found that Mongersen treatment rescued TZP number and density, follicle growth, and oocyte maturation, despite being cultured in a stiff microenvironment (Fig. 4e-j).

To understand how stromal stiffness ultimately affects the oocyte, we analyzed our RNA-seq data of isolated oocytes from cultured follicles. In the stiff hydrogel, GO terms of ‘mitochondrial gene expression’ and ‘NADH dehydrogenase complex assembly’ were downregulated (Fig. S3g), suggesting that mitochondrial function, including ATP production, may be impaired. Using tetramethylrhodamine methyl ester (TMRM) and MitoTracker Green (MTG), we indeed found reduced mitochondrial membrane potential and ATP production in germinal vesicle (GV)-stage oocytes from follicles cultured in stiff hydrogels (Fig. 5a, b), which is also consistent with isolated aged oocytes (Fig. S6a, b). Importantly, we found that Mongersen-treated follicles cultured in stiff hydrogels could restore mitochondrial function (Fig. 5a-c). Taken together, these data demonstrate that the restoration of stiffness-impaired TGF-β signaling via Mongersen partially rescues TZP, oocyte, and follicle dysfunction observed in stiff microenvironments characteristic of aged tissue mechanics.

**Figure 5.**
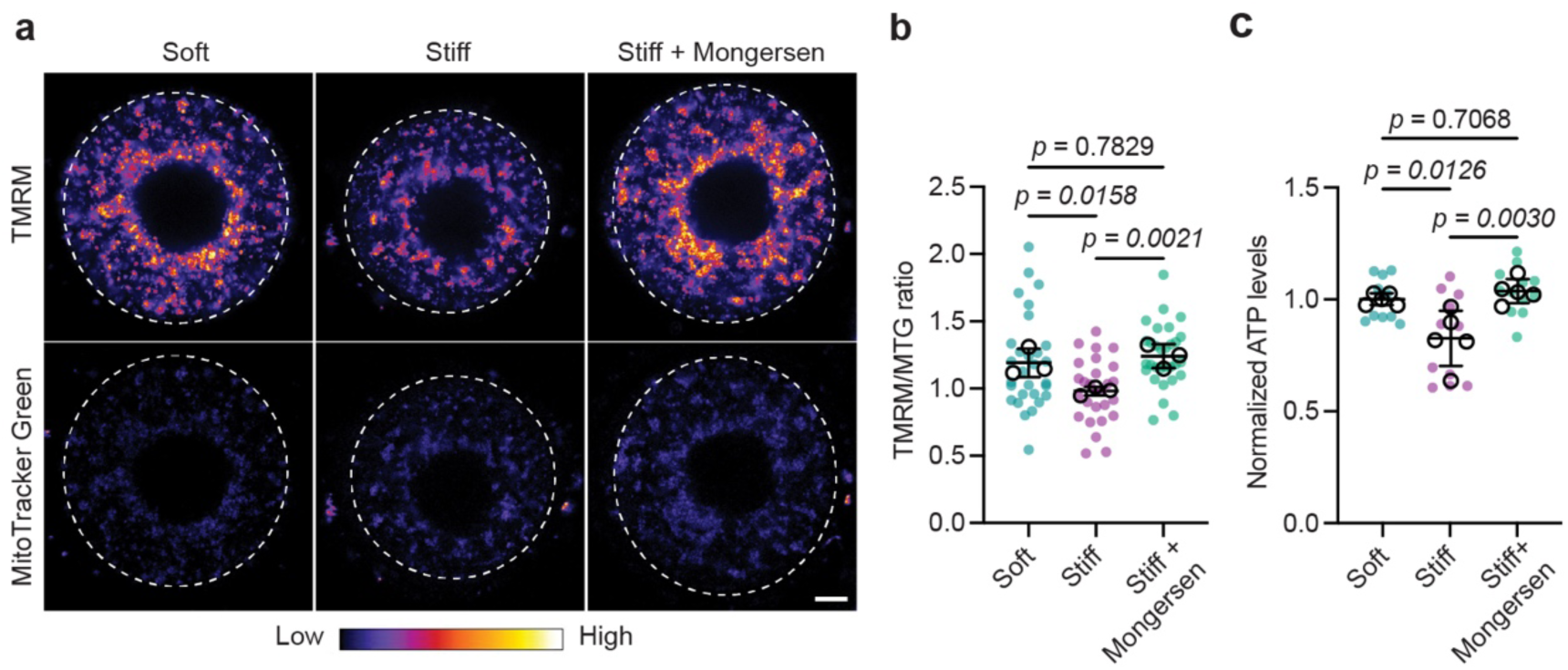
Targeting TGF-β signaling rescues oocyte mitochondrial defects in stiff hydrogels. Mitochondrial membrane potential was assessed through (a) tetramethylrhodamine methyl ester (TMRM) and MitoTracker Green (MTG) staining of live oocytes cultured in soft hydrogels, stiff hydrogels, and stiff hydrogels + Mongersen. (b) Mitochondrial potential (TMRM/MTG ratio) and (c) ATP levels were quantified for each group, showing an improvement with Mongersen treatment. (b,c) teal, light teal, and pink dots are individual oocytes; black circles represent biological replicates. Data is shown as mean ± SD and analyzed using One-way ANOVA. Sample size is: (b) n = 29, 27, and 30 in soft, stiff, and stiff + Mongersen groups, respectively. N = 3 replicates. (c) n = 10, 11, and 12 in soft, stiff, and stiff + Mongersen groups, respectively. N = 5 replicates. Scale bar: 30 µm.

## Discussion

In this study, we showed that age-associated stiffening of the ovarian stroma is sufficient to compromise follicle development and oocyte quality by disrupting soma-germLine communication. Through biochemical and biomechanical characterization of the aging ECM, combined with tunable hydrogels that recapitulate age-related stromal stiffness, we provide direct evidence that a stiff microenvironment alone can reduce follicular development and disrupt TZPs at the GC-oocyte interface. Importantly, we have identified TGF-β signaling as a mechanoregulated pathway linking matrix stiffness to impaired TZP formation. We further demonstrated that restoring this pathway can rescue follicular function despite a persistent stiff microenvironment.

Our results using spatial nanoindentation of ovarian tissue sections and atomic force microscopy on isolated follicles are consistent with recent follicular elastography measurements on whole ovaries (*27*). In young ovaries, follicles are almost two times stiffer than their surrounding stroma. However, ECM accumulation during aging causes a mechanical imbalance, with the stroma becoming stiffer than the follicles. Although the exact mechanisms underlying this age-related mechanical shift between follicles and stroma remain unclear, increased collagen production and reduced ECM turnover in ovarian stromal fibroblasts, coupled with minimal alterations to the ECM surrounding GCs, may contribute to this relationship (*40*). Consequently, aged follicles are exposed to higher mechanical stress compared to young follicles, and we found that this stiffer microenvironment restricts follicle activation and growth, consistent with previous studies showing that proper follicle development requires a soft matrix (*41*, *42*). It is important to note that aging-associated stromal ECM dysregulation not only alters stiffness, but also matrix composition and organization, and these cues likely cooperatively influence follicle function *in vivo*. For example, laminin, which increases in aged ovaries, could modulate follicle development via enhanced integrin-mediated adhesion formation (*43*, *44*). How cells integrate biochemical, architectural, and mechanical cues to regulate follicle function require further investigation using approaches such as bioengineered materials that can decouple contributions from these interconnected matrix signals in tissues (*45*).

Our results showed that a target of stromal stiffening is TZPs, structures that mediate GC-oocyte communication. Whereas intrinsic changes in GCs or oocytes have been shown to regulate TZPs (*5*, *46*, *47*), aging and mechanics can also play a role (*7*, *17*). Our data further demonstrated that TZP impairment due to higher ECM stiffness disrupts GC-oocyte connectivity. Such disruption could limit the transfer of essential metabolites, such as pyruvate and lactate, that support oocyte mitochondrial function (*48*). Oocyte maturation is an energy-intensive process, and the reduced ATP availability and mitochondrial membrane potential that we observed in stiff hydrogels could directly contribute to defective meiotic progression and abnormal spindle organization. Together, these findings support a model in which age-associated stromal stiffening disrupts TZPs and impairs soma-germLine communication during ovarian aging.

Our work further uncovered the TGF-β signaling pathway as a key link between stiff matrix and impaired follicle development. Although previous studies established TGF-β family growth factors such as GDF9 and BMP15 as essential regulators of TZP maintenance (*6*, *49*), our work advances this understanding by demonstrating that the mechanical microenvironment inhibits TGF-β signaling by upregulating Smad7, thereby suppressing Smad2/3 nuclear translocation. We showed that restoration of TGF-β activity via Smad7 knockdown using Mongersen was sufficient to rescue TZP density, follicle growth, and oocyte metabolic competence, suggesting that stiffness-induced follicle dysfunction is at least partially reversible. Our results implicate mechanoregulation of TGF-β signaling as a potential therapeutic target for improving follicle growth in aging or fibrotic ovarian environments, such as in polycystic ovary syndrome (PCOS) (*50–52*).

Future studies are required to understand how matrix stiffness is sensed by follicular cells leading to increased Smad7 expression (*53*, *54*). Inhibiting Smad7 only partially restored stiffness-mediated follicle and oocyte dysfunction, indicating that additional mechanisms may act in parallel. We also note that Smad7 inhibition in this study was carried out on young follicles cultured in stiff microenvironments. Therefore, further investigation is required to determine whether Smad7 inhibition in aged follicles will restore TZP density and follicle competence. Despite these open questions, our results establish stromal ECM stiffness as an important regulator of oocyte-follicle crosstalk and provides new insights into the mechanisms of follicle decline during female reproductive aging.

## Methods

### Animal and tissue handling

All animal experiments described here were approved by the Institutional Animal Care and Use Committee (IACUC) of the National University of Singapore (protocol R24-0684) and performed according to institutional guidelines. Reproductively young C57BL/6JInv(Jax) female mice (6-8 weeks old) were obtained from Singapore InVivos and housed at the National University of Singapore under standard laboratory conditions with free feeding, controlled temperature (21–25 °C) and humidity (30–70%), and maintained on a 12-hour light/dark cycle (lights on at 07:00). Reproductively aged mice were housed until 14-16 months old. Mice were kept virgin throughout the experimental period.

Ovarian tissues were collected immediately after euthanasia by carefully dissecting intact ovaries from the surrounding bursa under a stereomicroscope. Ovaries from both young and aged mice were isolated for downstream applications, including mass spectrometry, immunohistochemistry, nanoindentation of tissue sections, follicle AFM measurements, oocyte mitochondrial potential assays, and TZP immunostaining. Young ovaries were additionally used for *in vitro* follicle culture experiments.

### Alginate hydrogel synthesis

Alginate powder (Sigma-Aldrich, 71238) was dissolved in ultrapure water at 2% (w/v) by continuous stirring overnight in a 50 mL conical tube. Activated charcoal (Sigma-Aldrich) was added at a ratio of 0.5 g charcoal per 1 g alginate and mixed for 30 min. The suspension was filtered through a 0.22 µm filter (Millipore Express), frozen at −80 °C for 24 h and lyophilized for ∼1 week. Lyophilized alginate was stored at −80 °C until use. Prior to use in experiments, alginate was reconstituted in 1X DPBS lacking Ca²⁺ and Mg²⁺ to 2% (w/v), briefly vortexed, and placed on an orbital shaker until fully dissolved.

Hydrogel stiffness was tuned by varying CaCl₂ concentration and crosslinking duration. For soft hydrogels, 4.5 µL alginate droplets were crosslinked in 15 mM CaCl₂ and 140 mM NaCl for 30 s, then washed a single time in maintenance medium. For stiff hydrogels, droplets were first crosslinked in 15 mM CaCl₂ and 140 mM NaCl for 30 s, then transferred to 100 mM CaCl₂ for an additional 3 min and washed three times in maintenance medium.

### Follicle encapsulation and culture

Ovaries were isolated from young virgin C57BL/6J mice. Secondary follicles were then mechanically isolated in dissection medium consisting of Leibovitz’s L-15 (Thermo Fisher Scientific, 11415064) supplemented with 1% fetal bovine serum (FBS) and 100 U/mL penicillin-streptomycin. Isolated follicles were transferred to maintenance medium (αMEM; Thermo Fisher Scientific, 32561102) supplemented with 5% FBS and 100 U/mL penicillin-streptomycin for 2 h prior to encapsulation. After incubation, follicles with similar size were selected under a stereomicroscope. Individual follicles were encapsulated in 4.5 µL beads of 2% (w/v) alginate followed by a single wash in the maintenance medium. Encapsulated follicles were cultured individually in 96-well plates in 100 µL growth medium composed of a 1:1 mixture of αMEM GlutaMAX and F-12 GlutaMAX, supplemented with 5% FBS, 100 U/mL follicle-stimulating hormone (FSH; Sigma-Aldrich), and 5 µg/mL each of insulin, transferrin and selenium. Cultures were maintained at 37 °C and 5% CO₂, with 50% medium exchanged every 2 days for up to 7 days. Follicle diameter is tracked every 2 days.

### Mass spectrometry

Ovaries were collected from young and aged mice (n = 3, with each replicate consisting of ovaries from 4–5 mice). For each replicate, 8–10 ovaries were pooled to yield ∼20 mg fresh tissue. ECM enrichment was performed on the ovaries using the Compartmental Protein Extraction Kit (Merck Millipore), as previously described (*19*). Briefly, ovaries were flash-frozen in liquid nitrogen, pulverized with a mortar and pestle under liquid nitrogen, and resuspended in Lysis Buffer C (Merck Millipore) supplemented with 1X protease inhibitor (Merck Millipore)) at a final concentration of ∼20 mg tissue per 400 µL lysis buffer. Cytosolic, nuclear, membrane, and cytoskeletal fractions were sequentially extracted using the supplied buffers with centrifugation-based separation. The remaining ECM-enriched pellet was washed twice in Buffer W and three times in 1X PBS prior to preparation for mass spectrometry. All buffers were supplemented with 1X protease inhibitor (Merck Millipore), and Buffer N was supplemented with 3.5 µL Benzonase nuclease (Merck E1014) to digest genomic DNA and RNA.

Samples were resuspended in 100 μL of 8 M urea in 100 μM triethylammonium bicarbonate (TEAB) and lysed by sonication (15 s pulse followed by 45 s cooling, repeated three times). Proteins were precipitated by addition of four volumes of cold acetone and incubated at −80 °C for 2 d. Samples were centrifuged at 20,000 × g for 15 min at 4 °C, and the supernatant was removed. Pellets were resuspended in 50 μL of 8 M urea in 100 μM TEAB, and protein concentration was determined using the Pierce 660 nm protein assay. For reduction, 50 μg of protein from each sample was treated with 20 μM tris(2-carboxyethyl)phosphine (TCEP) in 100 μM TEAB for 30 min at room temperature. Alkylation was performed using chloroacetamide (CAA) to a final concentration of 55 μM in 100 mM TEAB for 30 min in the dark. Proteins were digested with Lys-C (1:25, enzyme:protein) for 4 h, followed by dilution to 1 M urea with 100 μM TEAB and overnight digestion with trypsin (1:50, enzyme:protein). Peptides were desalted using 1 mL C18 Empore SPE cartridges. For quantitative analysis, 5 μg of peptides per sample were labelled using TMT 6-plex reagents according to the manufacturer’s instructions. Labelled peptides were fractionated using high-pH reversed-phase fractionation prior to LC–MS/MS analysis. MS data were acquired on a Q Exactive HFX mass spectrometer (Thermo) using data-dependent mode.

### Atomic force microscopy (AFM)

Follicles were mechanically isolated from young and aged ovaries using dissection needles in Leibovitz’s L-15 medium (Thermo Fisher Scientific) supplemented with 1% FBS. Follicles with a spherical morphology and diameter ranging from 80 to 200 µm were selected. Surrounding follicular ECM was removed to minimize potential contributions of residual ECM to mechanical measurements. To immobilize follicles during indentation, a polydimethylsiloxane (PDMS) mold containing microwells was fabricated. Wells were 80 µm in depth and varied in diameter (100–300 µm) to accommodate different follicle sizes. Prior to use, PDMS molds were coated with poly-ʟ-lysine (0.1 mg/mL) for 10 min and washed twice with sterile water. Isolated follicles were transferred into wells of diameters slightly larger than the follicle diameter in L-15 medium. AFM indentation was performed using a NanoWizard 4 system (JPK Instruments/Bruker). A custom spherical probe was generated by adhering a 25 µm polystyrene sphere (Merck, 79807) onto a tip-less cantilever (Nanoworld ArrowTM TL1), yielding a soft cantilever with a spring constant of 0.03 N m⁻¹. Cantilevers were calibrated before each experiment using the thermal oscillation method in culture medium on glass. For each follicle, force measurements were obtained from four regions distributed across the apical surface. In each region, a 6 × 6 array of indentation was performed over a 10 × 10 µm² grid using Contact Mode. The maximum Vertical deflection setpoint was 8 nN, indentation depth is about 2 µm. Data were analyzed using the JPK analysis software after baseline correction. The Young’s modulus was calculated by fitting force indentation curves to the Hertz model (equation is shown below), where *F* is force, *E* is Young’s modulus, ν is Poisson’s ratio, *R* is the indenter radius, and *δ* is indentation depth. A Poisson’s ratio of 0.5 was assumed for all calculations.

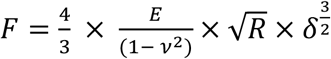

### Nanoindentation

Young and aged fresh ovaries were embedded in 5% (w/v) low-melting point agarose (Thermo Fisher Scientific) and sectioned into 80 µm-thick slices using a vibratome (Leica VT 1200S). Sections were released from the agarose and incubated with CNA35-GFP (100 µL) for 1 h at room temperature, followed by three washes in Leibovitz’s L-15 medium. For immobilization during indentation, glass-bottom dishes were coated with poly-D-lysine (0.1 mg/mL; Sigma-Aldrich, P1024). Ovary sections were transferred to coated dishes in L-15 medium supplemented with 1% FBS. Prior to nanoindentation, sections were imaged by epifluorescence microscope to quantify CNA35 signal using identical acquisition settings for all samples (4× objective, 5% excitation power and 40% gain). Mechanical measurements were performed using an Optics11 Life Chiaro Nanoindenter equipped with a spherical probe (nominal spring constant, 0.02 N/m; sphere diameter, 50 µm). To minimize sample-probe adhesion, the probe was coated with 1% Pluronic (Sigma-Aldrich) before measurements. Probe positioning was guided by wide-field microscopy, and measurements were targeted to ovarian stroma. Quasi-static matrix scans were acquired with a 2 µm indentation depth using a 4 × 4 grid with a 10 µm step size. Young’s modulus at each location was obtained by fitting force-indentation curves to the Hertz contact model in the Optics11 DataViewer software with Poisson’s ratio ν = 0.5. For each group, n = 3-5 mice were analyzed, with one tissue section collected from each ovary. To correlate tissue stiffness with ECM abundance, two images were acquired from the same ovary section: an epifluorescence image of CNA35-labelled collagen and a wide-field image showing the nanoindenter probe positioned on the tissue surface. The wide-field image was imported into Adobe Illustrator, adjusted to 50% opacity, and then rotated and scaled to match the fluorescence image. The two images were subsequently overlaid to spatially map the indentation site onto the corresponding region of the fluorescence image.

For alginate hydrogels, soft and stiff hydrogel beads were immobilized on glass dishes using superglue, and stiffness was measured using the same cantilever and indentation settings unless noted. Quasi-static scans were acquired at a 2 µm indentation depth using a 2 × 2 grid with 10 µm step size. Young’s modulus was calculated using the same analysis pipeline. A total of n = 10 soft and n = 28 stiff hydrogel beads were measured.

### RNA sequencing

Follicles were cultured in soft or stiff hydrogels for 6 days, retrieved from the gels and washed twice in maintenance medium. Cumulus GCs were removed and oocytes were denuded by gentle aspiration using a fine mouth pipette, followed by a 1 min digestion with 0.25% trypsin. Denuded oocytes were immediately transferred into lysis buffer (NEB) and stored at −80 °C until reverse transcription. Five oocytes were pooled per sample. GC niches were collected for RNA isolation using the RNeasy Plus Micro Kit (Qiagen) according to the manufacturer’s instructions. Briefly, GCs were transferred into 350 µL RLT buffer and homogenized by pipetting. Ethanol (250 µL; absolute) was added to the lysate, mixed thoroughly, and the mixture was loaded onto a RNeasy MinElute spin column placed in a 2 mL collection tube. Columns were centrifuged for 15 s at 8,000 × g, washed with 500 µL RPE buffer (15 s at 8,000 × g), followed by a wash with 500 µL 80% ethanol (2 min at 8,000 × g). Columns were then placed in a new 2 mL collection tube and dried by centrifugation at maximum speed for 5 min. RNA was eluted in 14 µL RNase-free water. RNA quality was assessed using TapeStation High Sensitivity RNA ScreenTape (Agilent), and samples with an RNA integrity number equivalent (RINe) > 8.5 were processed further. cDNA synthesis and sequencing library preparation were performed using the NEBNext Single Cell/Low Input RNA Library Prep Kit for Illumina according to the manufacturer’s instructions. For each sample, 20 ng total RNA was used for cDNA synthesis and amplified for 8 PCR cycles. Libraries were indexed using NEBNext Multiplex Oligos for Illumina and sequenced by NovogeneAIT Genomics (Singapore) on an Illumina NovaSeq 6000 platform.

### Immunostaining

Ovarian tissues and isolated follicles were fixed in 4% paraformaldehyde (PFA) for 40 min and washed three times in PBS. For immunohistochemistry (IHC), ovaries were embedded in OCT and cryosectioned at 10 µm using a cryostat (Leica CM1950) onto Superfrost slides, while follicles were embedded in 5% (w/v) agarose and sectioned at 50 µm using a vibratome (Leica VT1200) and keep in the PBS. For oocyte immunofluorescence (IF), isolated oocytes were fixed in pre-warmed (37 °C) 4% PFA for 20 min and washed three times in 1X PBS within a hydrophobic barrier. All samples were permeabilized with 0.5% Triton X-100 for 20 min, blocked with 5% BSA for 2 h, and incubated with primary antibodies overnight at 4 °C, followed by incubation with appropriate secondary antibodies for 1 h at room temperature. Primary antibodies used were rabbit anti-collagen IV (Abcam, ab236640; 1:150), rabbit anti-collagen I (Abcam, ab21286; 1:150), rabbit anti-laminin (Invitrogen, PA1-16730; 1:150), rabbit anti-Ki-67 (D3B5) (Cell Signaling Technology, #9129; 1:200), mouse anti-α-tubulin (Sigma, F2168; 1:500), rabbit anti-YAP1 (Cell Signaling Technology, #14074; 1:200), rabbit anti-SMAD2/3 (D7G7) (Cell Signaling Technology, #8878; 1:150), and plasmid CNA35 for collagen (Addgene, #61603; 1:100). For secondary antibodies, Alexa Fluor 647-conjugated goat anti-rabbit IgG (Invitrogen, SA5-10395-AF647; 1:1000) was used. Alexa Fluor® 488 Phalloidin (Cell Signaling Technology, #8686; 1:400) was used to stain F-actin, and Hoechst 33342 (Invitrogen, H3570; 1:1000) was used for nuclear staining. Ovary ECM IHC images were acquired on an Olympus FV3000, oocyte and follicle IF samples were imaged using Zeiss LSM 980 laser-scanning confocal microscope.

### Mitochondrial potential staining

Germinal vesicle (GV) stage oocytes were isolated from follicles and denuded of surrounding GCs. Oocytes were incubated in growth medium containing 25 nM tetramethylrhodamine methyl ester (TMRM; Invitrogen, T668) and 100 nM MitoTracker Green (MTG; Invitrogen, M7514) for 30 min at 37 °C and 5% CO_2_, protected from light. Oocytes were then washed 3 times and transferred to fresh culture medium for live imaging. TMRM and MTG fluorescence were acquired using a Zeiss LSM 980 scanning laser confocal microscope under identical settings across groups.

### Mongersen treatment

Mongersen (MedChemExpress, HY-145721) was dissolved in water to prepare a 1 mM stock solution and stored at −80 °C. Mongersen was added to follicle culture medium at a final concentration of 0.5 µM on day 0. After 2 days of culture, the concentration was reduced to 0.1 µM by complete medium replacement with fresh growth medium containing Mongersen at 0.1 µM, and 0.1 µM was maintained until the experimental endpoint. Control follicles were treated with an equivalent volume of water (vehicle). Mongersen concentrations were empirically titrated, as higher concentrations inhibited follicle growth. Quantification of follicle growth rate, TZP number and density, Smad2/3 nuclear-to-cytoplasmic ratio, oocyte maturation rate, mitochondrial membrane potential, and oocyte ATP levels for Mongersen-treated follicles was performed using the same analysis methods detailed in the corresponding methodological sections.

### Oocyte maturation

Follicular structures were released from alginate beads using 10 IU/mL alginate lyase (Sigma-Aldrich, A1603). Cumulus-oocyte complexes were transferred to maturation medium consisting of αMEM supplemented with 10% FBS, 1.5 IU/mL human chorionic gonadotropin, 10 ng/mL epidermal growth factor, and 100 mIU/mL follicle-stimulating hormone. *In vitro* maturation was carried out for 16 h at 37 °C in 5% CO₂. Metaphase II (MII) oocytes were identified by the presence of the first polar body extrusion under the stereomicroscope. Oocytes matured *in vitro* were then fixed and stained for spindle and chromosomes as described in the immunostaining section.

### Oocyte ATP measurements

Total ATP content was measured in pools of three oocytes using the ATP Bioluminescent Somatic Cell Assay Kit (Sigma-Aldrich, FLASC) according to the manufacturer’s instructions. Briefly, the ATP assay working solution (100 µL; 1:25 dilution of stock) was dispensed into each well of a 96-well plate and equilibrated for 3 min at room temperature. Samples were prepared by mixing 50 µL oocyte lysate with 100 µL somatic cell ATP releasing reagent and 50 µL filtered ultrapure water. A 100 µL aliquot of each sample mixture was transferred to assay wells, and luminescence was measured immediately using a microplate reader (Varioskan LUX, Thermo Fisher Scientific).

### Image analysis and quantification

Follicle diameter was measured in FIJI using the line tool by recording the major and minor axes of each follicle at day 0 and day 6. The mean diameter was defined as the average of the major and minor axis lengths. Follicle growth rate was calculated as Growth Rate = (Diameter at day 6 – Diameter at day 0) / 6 days.

All oocyte spindle images were processed using ZEN 3.8, and three-dimensional reconstructions of tubulin and DNA channels were generated from z-stacks encompassing the entire spindle. Spindles were classified as normal if they exhibited a compact bipolar structure with focused poles and a clear axis, whereas abnormal spindles showed one or more of the following features: multipolarity (≥3 poles), unfocused or broadened poles, disorganized microtubules (splayed, fragmented, or irregular bundles with loss of bipolar architecture), or bent/asymmetric morphology. Chromosome alignment was evaluated on the DNA channel. Chromosomes were considered aligned when a compact metaphase plate at the spindle equator was formed, and misaligned when at least one chromosome was located outside the metaphase plate or dispersed along the spindle axis (*55*). The spindle abnormality rate was calculated as the percentage of abnormal oocytes among the total analyzed.

Ovary ECM IHC images were analyzed using FIJI (ImageJ). Images were acquired under identical exposure settings across groups. Fluorescence intensity of ECM markers (collagen I, collagen IV, and laminin) was quantified from manually defined regions of interest (ROIs) corresponding to follicles and stromal areas. ROIs were delineated based on the phalloidin channel to define follicular architecture. Fluorescence intensity was normalized to ROI area and expressed as the mean gray value. Sections processed in parallel without primary antibodies served as background controls, and background-corrected intensities were calculated by subtracting the mean control value from each corresponding measurement. All analyses were performed using identical processing parameters. Mean CNA35 intensity was quantified as the mean gray value within the ovary tissue region. High collagen was assessed by calculating the fraction of tissue area exhibiting CNA35 signal intensities above 1000, expressed relative to the total area of the ovary section.

Ki-67-positive nuclei were quantified in FIJI using the DAPI channel to define nuclei. For ovary slices, follicle regions were selected with polygon selection tool and copied. Nuclei were segmented from the DAPI channel by 8-bit conversion, global thresholding (Otsu), binarization and watershed separation, and counted using Analyze Particles with identical size and circularity parameters applied across all images. Nuclear ROIs derived from the DAPI mask were overlaid onto the Ki-67 channel to measure mean nuclear Ki-67 intensity. Ki-67-positive nuclei were defined using a fixed intensity threshold determined from background controls and applied consistently across samples. The percentage of Ki-67-positive nuclei was calculated as (Ki-67-positive nuclei / total DAPI nuclei) × 100.

YAP and Smad2/3 nuclear-to-cytoplasmic (N/C) ratios were quantified in FIJI. ROIs corresponding to follicular GCs were manually defined by excluding the oocyte and pixels outside the follicle. Total GC fluorescence intensity was measured for YAP or Smad2/3 within the GC ROI. Nuclei were segmented from the DAPI channel to generate a nuclear mask, which was applied to the YAP or Smad2/3 channel to quantify nuclear fluorescence intensity. Background intensity was measured in tissue-free regions and subtracted from both nuclear and whole-cell measurements. For each follicle, the N/C ratio was calculated as nuclear intensity / (whole-cell intensity − nuclear intensity).

TZP density was quantified from phalloidin-stained images using FIJI. Analysis was performed on a single confocal optical section at the oocyte equatorial plane. Four 10 µm arc segments, spaced 90° apart along the oocyte circumference, were defined, and TZPs within each arc were quantified by counting peaks in fluorescence intensity (Plot Profile + Find Peaks). TZP density was calculated as the mean number of TZPs per 10 µm arc across the four segments. Oocyte diameter and perimeter were measured to estimate total TZP number per oocyte. The number of GCs surrounding each oocyte was counted manually.

For live oocyte mitochondrial potential assessment, an ROI encompassing the entire oocyte was defined and mean fluorescence intensity per unit area was quantified for TMRM and MTG. Background intensity was measured in cell-free regions and subtracted. Mitochondrial membrane potential was quantified as the ratio of mean TMRM intensity to mean MTG intensity (TMRM/MTG).

### Bioinformatics analysis

Proteins were identified using Proteome Discoverer v2.3 (Thermo). Peptide- and protein-level identifications were filtered at a false discovery rate (FDR) ≤ 1% and quantified based on TMT reporter ion intensities. Protein identifiers were mapped against the UniProt reference proteome (*Mus musculus*). Normalization was performed using variance stabilizing normalization (VSN). Differentially abundant proteins were analyzed using the DEP R package (*56*). Matrisome proteins were annotated using the Matrisome Database (*21*, *57*). Data visualization was performed using GraphPad Prism 9 and TBtools (*58*).

For RNA-seq, differential gene expression analysis was performed using DESeq2 (Bioconductor 3.17). Gene annotation was based on Ensembl. Gene ontology (GO) enrichment analysis was performed using the WEB-based GEne SeT AnaLysis Toolkit (WebGestalt). Protein-protein interaction (PPI) analysis was performed using STRING. Heatmaps were generated using TBtools, volcano plots were generated using OriginPro 2024, and gene expression plots were generated using GraphPad Prism 9.

### Statistical analysis and sample sizes

No statistical methods were used to predetermine sample sizes; however, sample sizes were comparable to those in previous studies (*9*, *16*, *44*). Statistical analyses were performed using GraphPad Prism 9 and/or R. For comparisons between two groups, two-tailed unpaired Student’s t-tests were used. For comparisons among three groups, one-way analysis of variance (ANOVA) followed by Tukey’s multiple-comparison test was used. Exact P values are reported on graphs. Sample sizes (n) and the number of independent experiments are reported in the corresponding figure legends. Data are presented as mean ± s.d.. Unless otherwise stated, experiments were performed with at least three independent biological replicates (N ≥ 3). Data collection and analysis were not performed in a blinded manner. No data were excluded from the analyses, and no specific methods were used for random allocation of samples to experimental groups.

## Data availability

The mass spectrometry proteomics data will be deposited to the ProteomeXchange Consortium via the PRIDE repository with the dataset identifier TBD. The RNA-seq data will be deposited to NCBI GEO repository with accession code TBD.

## Acknowledgements

The authors thank Jennifer Marlena (MBI, NUS) for illustrations; Chii Jou Chan (MBI, NUS) for providing vibratome for usage; Lim Chwee Teck and Brenda Nai (MBI, NUS) for providing and assisting AFM usage; Mona Suryana and Foo Sebastian Junwei (MBI Nano and Microfabrication Core) for PDMS mold fabrication; and Hui Ting Ong (MBI, NUS) for image analysis support. We also thank Chii Jou Chan, Yan Jie, Hu Jinrong, and Wong Bin Sheng (MBI, NUS) for fruitful discussions and technical advice, as well as Hannah Fung for draft revision. We thank the MBI Wet Lab Core for facility and technical support, the Singapore Microscopy and Bioimaging Analysis (SiMBA, MBI) core for microscopy and data processing facilities, and the High-throughput Molecular Genetics (HMG, MBI) core for RNA-seq support.

This work was supported by the Bia-Echo Asian Centre for Reproductive Longevity and Equality (ACRLE) under the NUS Yong Loo Lin School of Medicine and the National Research Foundation, Singapore, under its Mid-sized Grant NRF-MSG-2023-0001, as well as the Ministry of Education under Start-up Grants in the Research Centres of Excellence programme through the Mechanobiology Institute at the National University of Singapore and the Biomedical Engineering Department at the National University of Singapore.

## Author contributions statement

J.L.Y., R.L., H.W., and X.S. conceptualized the project. J.L.Y., R.L., and X.S. wrote the manuscript. J.L.Y. and R.L. supervised data collection and analysis. X.S. performed most experiments and data analysis. H.W., G.C., and Y.L. assisted in experiments and data analysis. J.Z. and Y.L. assisted in RNA-seq experiments and analysis. L.C.W., T.Z., S.G.L., R.M.S. assisted in MS experiments and analysis.

## Competing interest statement

The authors declare there are no competing interests.

## Figure Legends

**Fig. S1.**
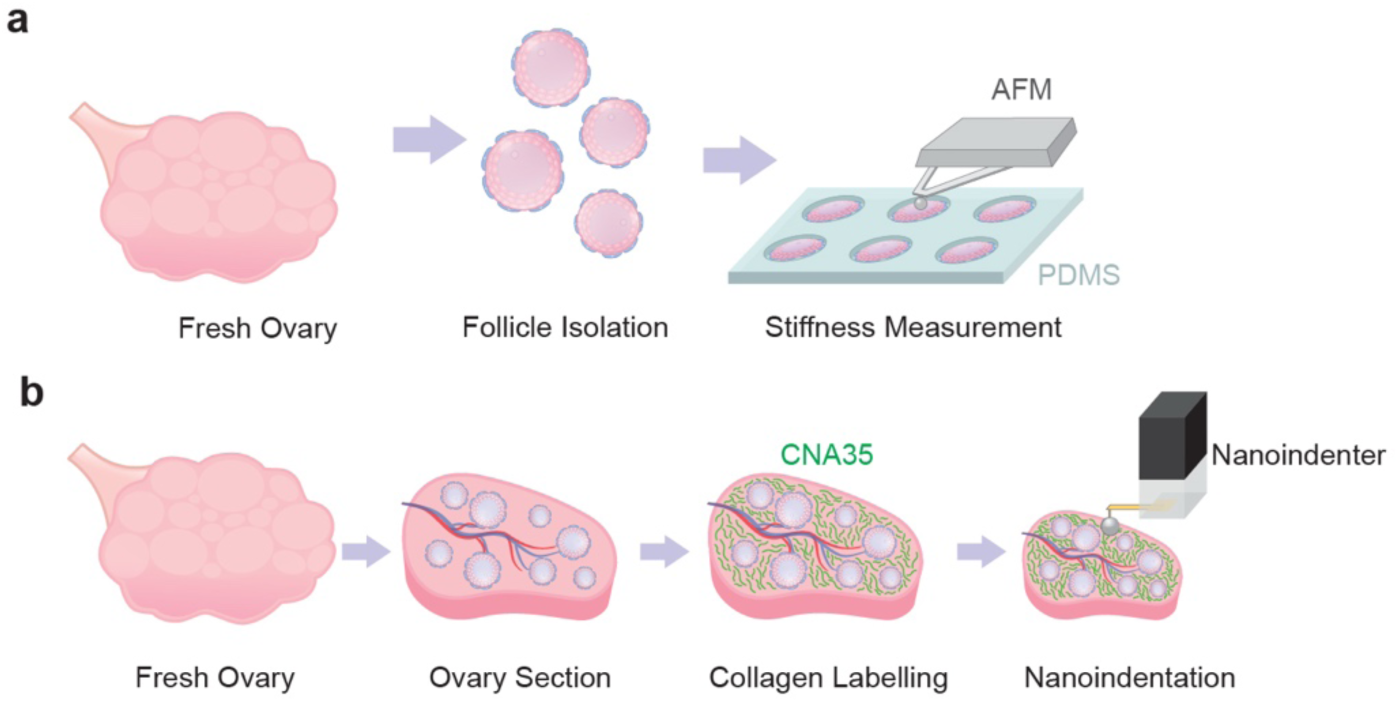
Mechanical measurements on follicles and ovarian stroma. (a) Follicle stiffness was measured using atomic force microscopy (AFM). Individual follicles were isolated from fresh ovaries (young and aged mice) and immobilized within a PDMS microwell array prior to AFM indentation. (b) Correlation of ovarian stroma ECM to stiffness was carried out by sectioning fresh ovaries and live-labeling with CNA35 (green) to visualize collagen, followed by stiffness measurements using a nanoindenter. Captured fluorescent images of collagen were correlated to the regions of indentation.

**Fig. S2.**
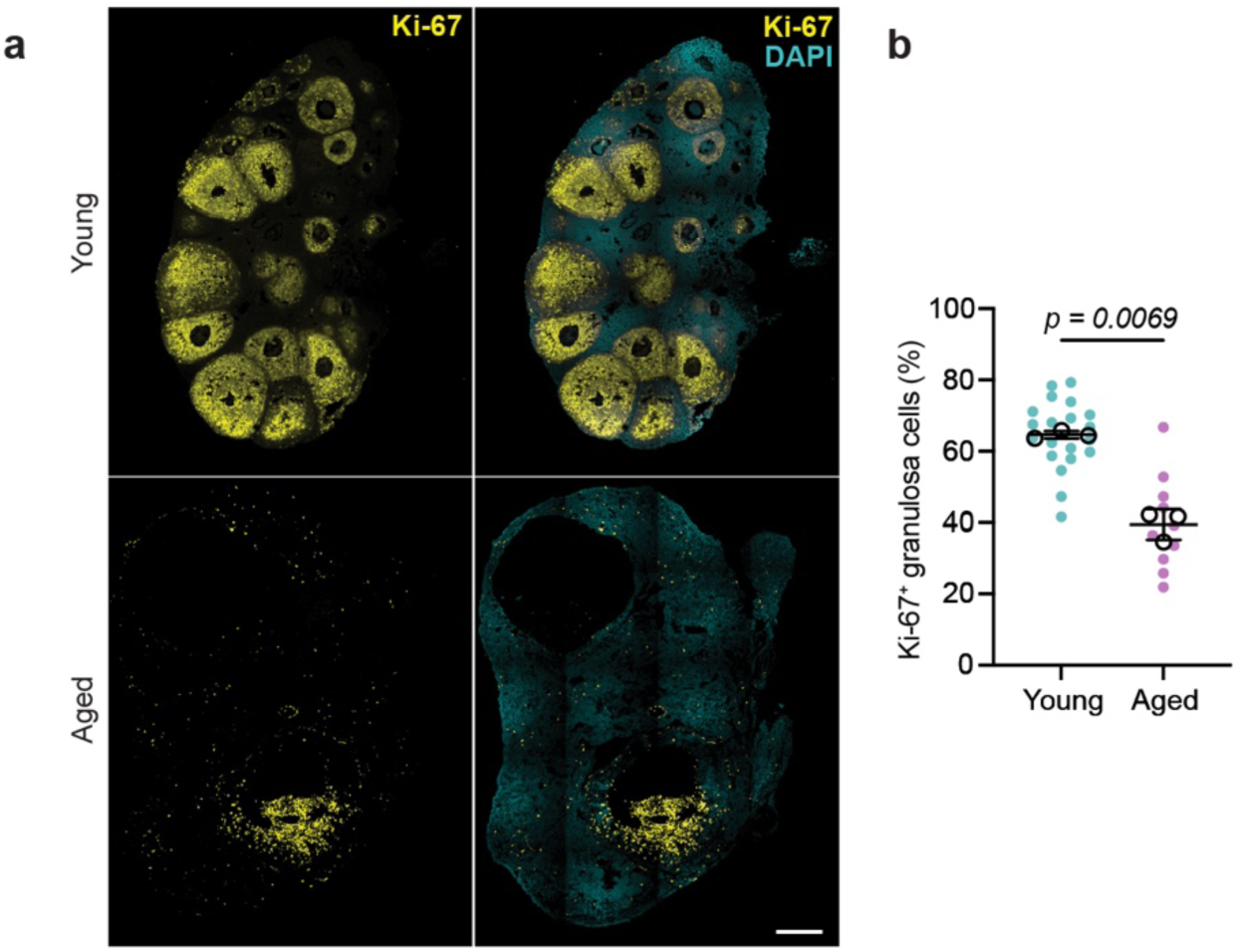
Cell proliferation declines in aged ovarian tissue. (a) Immunostaining of Ki-67 (yellow) and DAPI (cyan) in ovarian tissue cryosections from young (6 weeks) and aged (14 months) mice. (b) Quantification of cell proliferation, shown as the percentage of Ki-67 positive granulosa cells / follicle in young and aged groups. Data are presented as mean ± SD. n = 20 follicles in the young group and n = 9 follicles in the aged group. (b) teal and pink dots are individual follicles; black circles represent biological replicates. Statistical significance was determined by unpaired two-tailed Student’s t-test. Scale bar: 200 µm.

**Fig. S3.**
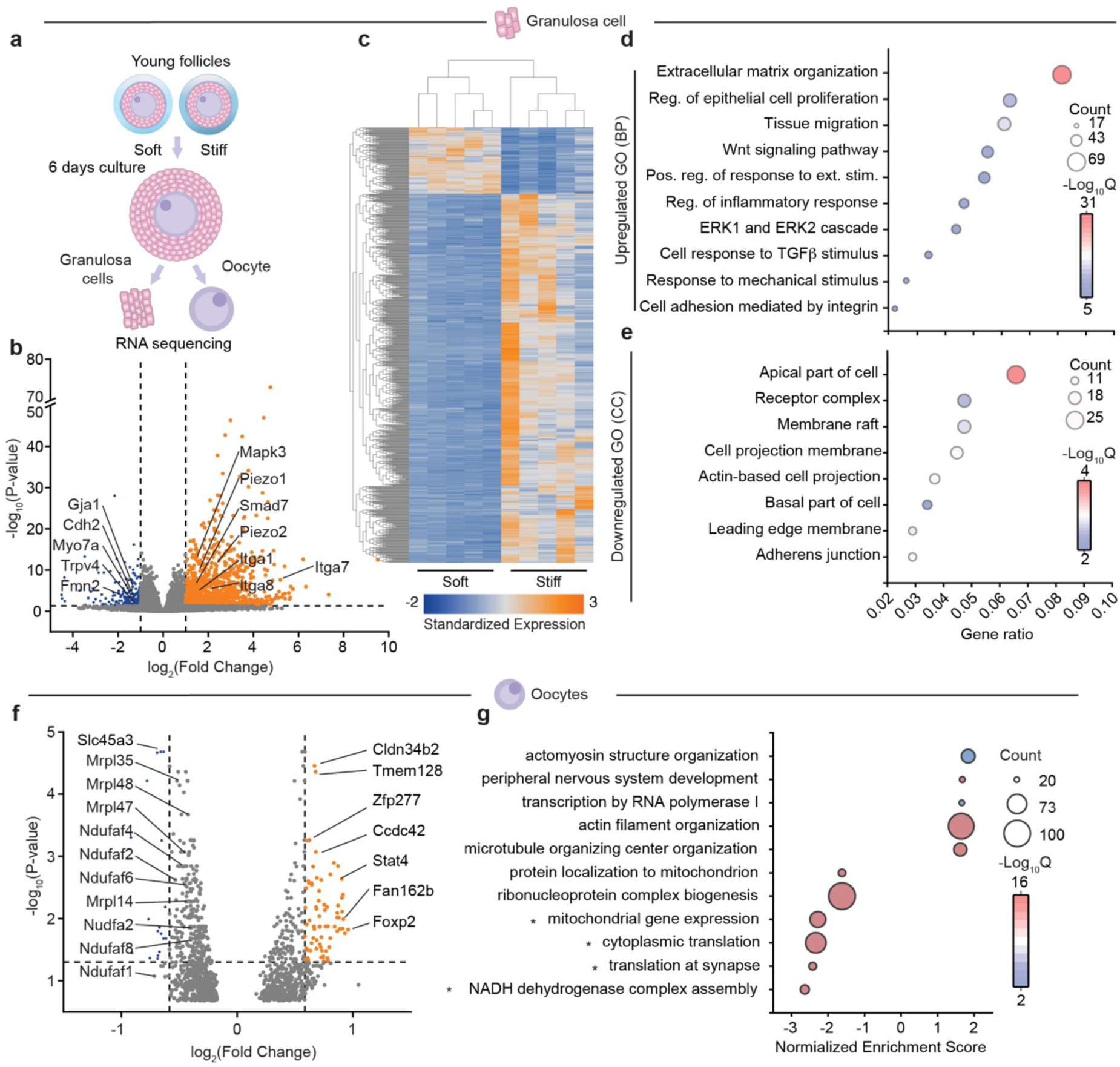
Differential gene expression of granulosa cells and oocytes cultured in soft versus stiff hydrogels. (a) After young follicles were cultured for 6 days in soft or stiff hydrogels, granulosa cells and oocytes were separately isolated for bulk RNA sequencing (RNA-seq). (b- e) Represent data from granulosa cell RNA-seq, while (f-g) represents oocyte RNA-seq. (b) Volcano plot showing DEGs in granulosa cells cultured in stiff vs. soft hydrogels with selected mechanoregulated genes highlighted. (c) Heatmap illustrating the hierarchical clustering of DEGs in granulosa cells from follicle culture in soft and stiff hydrogels. (d, e) GO enrichment analysis of downregulated cellular components (CC) and upregulated biological processes (BP) in stiff versus soft hydrogels. For oocytes, (f) mitochondrial-related genes are highlighted in the volcano plot of DEGs. (g) GSEA of top enriched pathways in stiff versus soft hydrogels showing. Graph shows Normalized Enrichment Scores (NES), with bubble size representing gene count and color indicating statistical significance (-Log_10_Q).

**Fig. S4.**
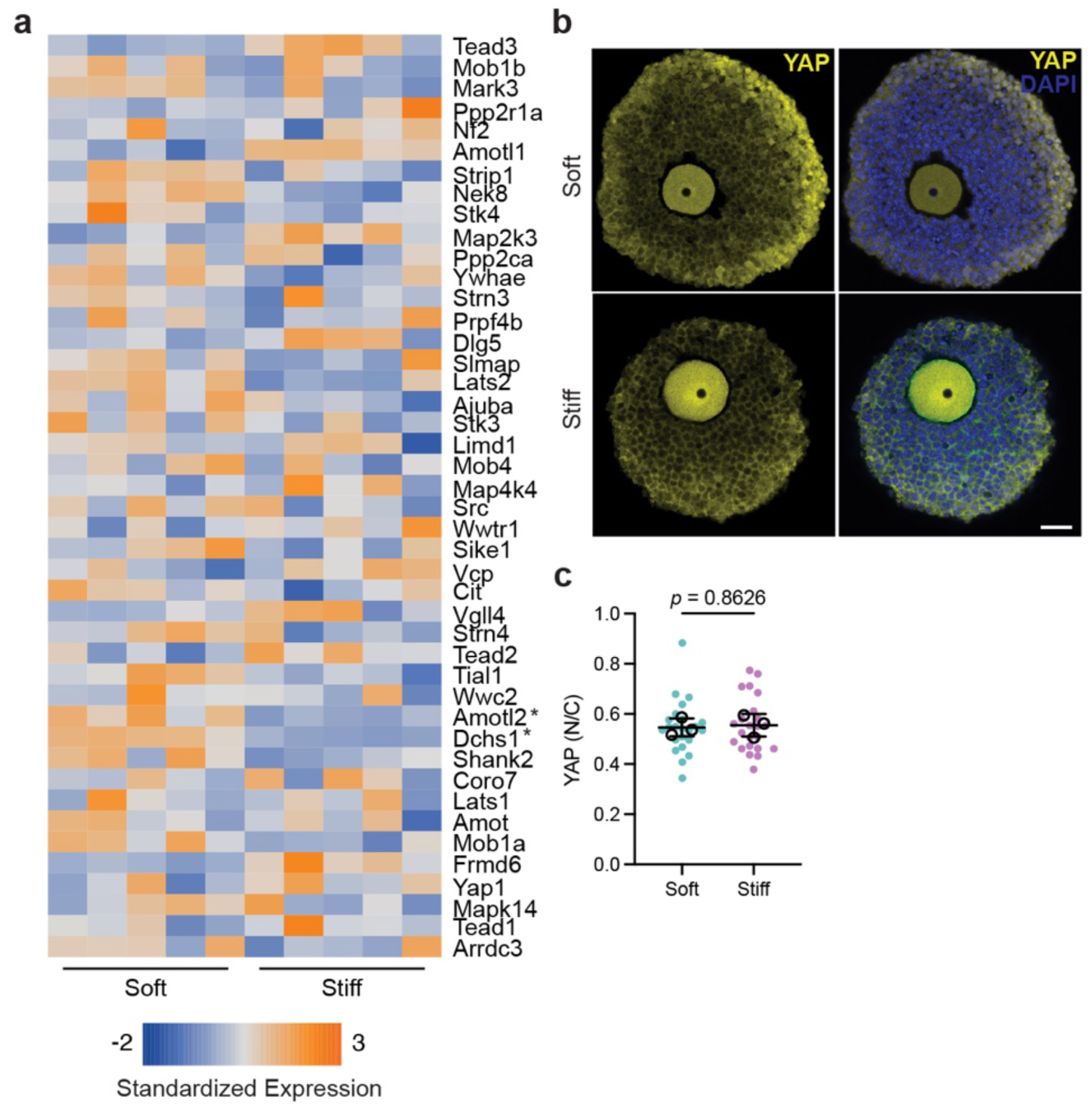
Hippo pathway analysis in granulosa cell RNA-seq data. (a) Heatmap shows Hippo pathway gene expression of granulosa cells isolated from follicles cultured in soft and stiff hydrogels. Asterisk (*) represents significance. At the protein level, (b) YAP (yellow) immunostaining staining in young follicles cultured in soft and stiff hydrogels was carried out with corresponding (c) nuclear (DAPI, blue) translocation quantification (nuclear to cytoplasmic fluorescence intensity ratio: N/C). (c) teal and pink dots are individual follicles; black circles represent biological replicates. Data is shown as mean ± SD and analyzed using unpaired t-test. Sample size is: 23 follicles from 3 independent experiments in both soft and stiff. Scale bar: 30 µm.

**Fig. S5.**
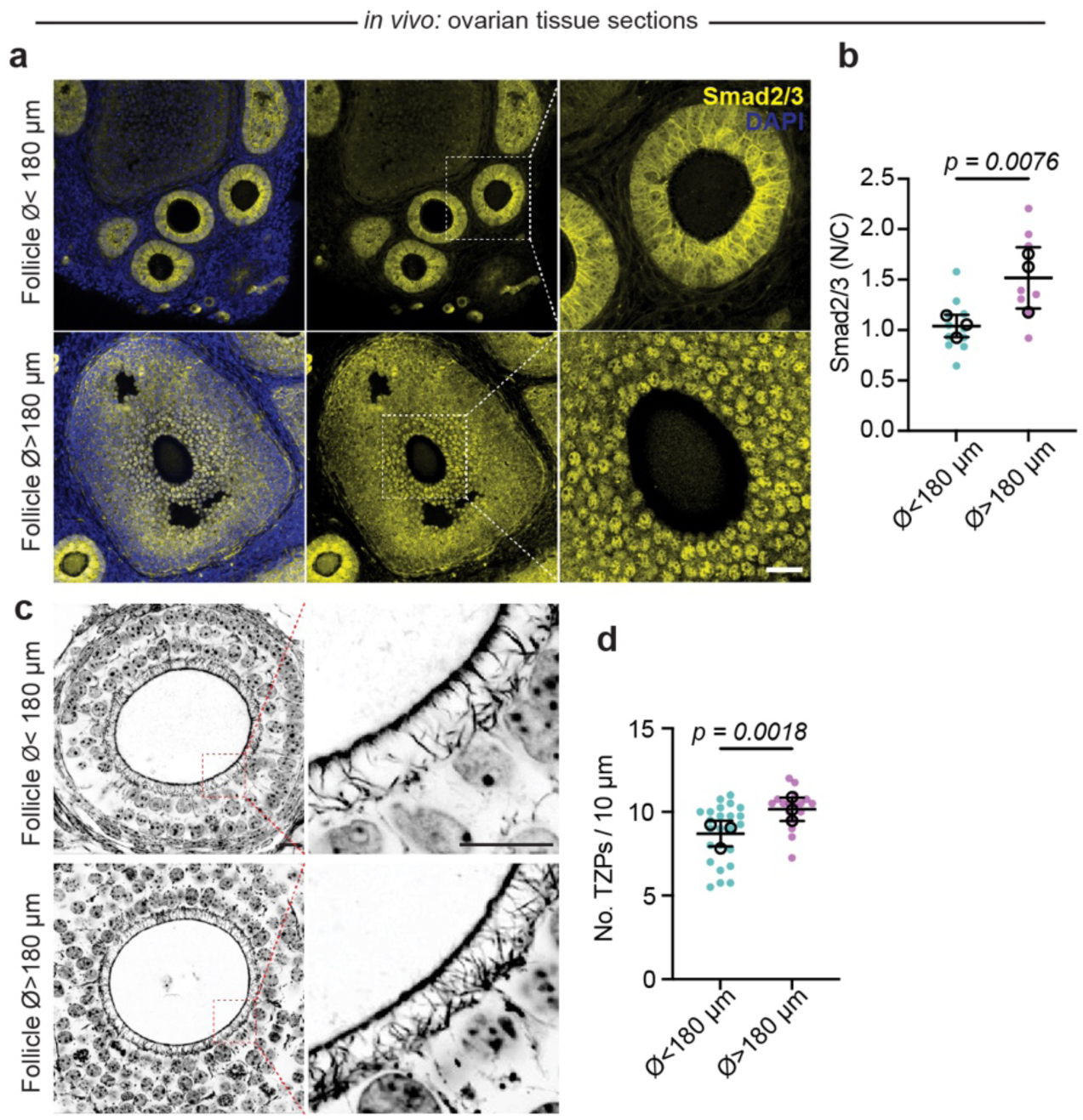
*In vivo* Smad2/3 localization of small and large follicles in ovarian tissue. (a) SMAD2/3 (yellow) and DAPI (blue) in young mice as a function of follicle size (follicle diameter ⌀ <180 µm - top versus ⌀ >180 µm - bottom). (b) Nuclear translocation of Smad2/3 in granulosa cells was quantified as a function of follicle diameter (nuclear to cytoplasmic fluorescence intensity ratio: N/C). (c) Actin immunostaining shows TZPs from (follicle diameter ⌀ <180 µm - top versus ⌀ >180 µm - bottom) in young mice. (d) TZP density as a function of follicle diameter was quantified. (b,d) teal and pink dots are individual follicles; black circles represent biological replicates. Data in (b) and (d) are presented as mean ± SD. Sample size is: (b) n = 11 (D < 180) and 7 (D > 180) follicles from 3 mice. (d) n = 25 (D < 180) and 18 (D > 180) follicles from 3 young mice. Statistical significance was determined by unpaired two-tailed Student’s t-test. Scale bar: 80 µm in (a) and 10 µm in (c).

**Fig. S6.**
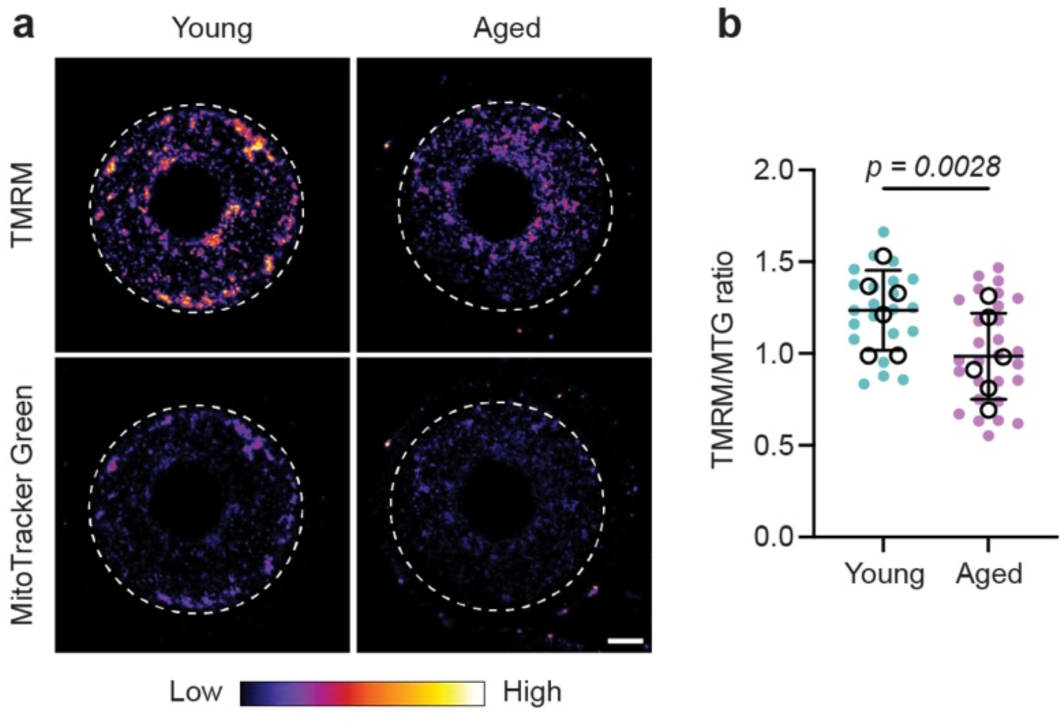
Mitochondrial membrane potential is reduced in aged oocytes. Mitochondrial membrane potential was assessed through (a) TMRM (top) and MTG (bottom) in freshly isolated young and aged oocytes. (b) Mitochondrial potential (TMRM/MTG ratio) was quantified, showing a reduction with age. b) teal and pink dots are individual oocytes; black circles represent biological replicates. Data is shown as mean ± SD and analyzed using unpaired t-test. Sample size : n = 22 (Young) and 25 (Aged) oocytes from 6 mice in each group. Scale bar: 20 µm.

## Notes

### Competing Interest Statement

The authors have declared no competing interest.

## References

1. L. J. Heffner, Advanced Maternal Age — How Old Is Too Old? N. Engl. J. Med. 351, 1927–1929 (2004).

2. J. R. Gruhn, A. P. Zielinska, V. Shukla, R. Blanshard, A. Capalbo, D. Cimadomo, D. Nikiforov, A. C.-H. Chan, L. J. Newnham, I. Vogel, C. Scarica, M. Krapchev, D. Taylor, S. G. Kristensen, J. Cheng, E. Ernst, A.-M. B. Bjørn, L. B. Colmorn, M. Blayney, K. Elder, J. Liss, G. Hartshorne, M. L. Grøndahl, L. Rienzi, F. Ubaldi, R. McCoy, K. Lukaszuk, C. Y. Andersen, M. Schuh, E. R. Hoffmann, Chromosome errors in human eggs shape natural fertility over reproductive life span. Science 365, 1466–1469 (2019).

3. D. Cimadomo, G. Fabozzi, A. Vaiarelli, N. Ubaldi, F. M. Ubaldi, L. Rienzi, Impact of Maternal Age on Oocyte and Embryo Competence. Front Endocrinol 9, 327 (2018).

4. A. Camaioni, M. A. Ucci, L. Campagnolo, M. D. Felici, F. G. Klinger, O. behalf of the I. S. of E. (SIERR) Reproduction and Research, The process of ovarian aging: it is not just about oocytes and granulosa cells. J Assist Reprod Gen 39, 783–792 (2022).

5. H. Wang, Z. Huang, X. Shen, Y. Lee, X. Song, C. Shu, L. H. Wu, L. S. Pakkiri, P. L. Lim, X. Zhang, C. L. Drum, J. Zhu, R. Li, Rejuvenation of aged oocyte through exposure to young follicular microenvironment. *Nat*. Aging, 1–17 (2024).

6. M. J. Carabatsos, J. Elvin, M. M. Matzuk, D. F. Albertini, Characterization of Oocyte and Follicle Development in Growth Differentiation Factor-9-Deficient Mice. Dev. Biol. 204, 373–384 (1998).

7. S. El-Hayek, Q. Yang, L. Abbassi, G. FitzHarris, H. J. Clarke, Mammalian Oocytes Locally Remodel Follicular Architecture to Provide the Foundation for GermLine-Soma Communication. Curr Biol 28, 1124–1131.e3 (2018).

8. C. Wu, D. Chen, M. B. Stout, M. Wu, S. Wang, Hallmarks of ovarian aging. Trends Endocrinol. Metab. 36, 418–439 (2025).

9. F. Amargant, S. L. Manuel, Q. Tu, W. S. Parkes, F. Rivas, L. T. Zhou, J. E. Rowley, C. E. Villanueva, J. E. Hornick, G. S. Shekhawat, J. J. Wei, M. E. Pavone, A. R. Hall, M. T. Pritchard, F. E. Duncan, Ovarian stiffness increases with age in the mammalian ovary and depends on collagen and hyaluronan matrices. Aging Cell 19, e13259 (2020).

10. C. W. McCloskey, D. P. Cook, B. S. Kelly, F. Azzi, C. H. Allen, A. Forsyth, J. Upham, K. J. Rayner, D. A. Gray, R. W. Boyd, S. Murugkar, B. Lo, D. Trudel, M. K. Senterman, B. C. Vanderhyden, Metformin Abrogates Age-Associated Ovarian Fibrosis. Clin Cancer Res 26, 632–642 (2020).

11. S. S. Dipali, C. D. King, J. P. Rose, J. E. Burdette, J. Campisi, B. Schilling, F. E. Duncan, Proteomic quantification of native and ECM-enriched mouse ovaries reveals an age- dependent fibro-inflammatory signature. Aging 15, 10821–10855 (2023).

12. S. M. Briley, S. Jasti, J. M. McCracken, J. E. Hornick, B. Fegley, M. T. Pritchard, F. E. Duncan, Reproductive age-associated fibrosis in the stroma of the mammalian ovary. Reproduction 152, 245–260 (2016).

13. D. A. Landry, E. Yakubovich, D. P. Cook, S. Fasih, J. Upham, B. C. Vanderhyden, Metformin prevents age-associated ovarian fibrosis by modulating the immune landscape in female mice. Sci Adv 8, eabq1475 (2022).

14. T. Umehara, Y. E. Winstanley, E. Andreas, A. Morimoto, E. J. Williams, K. M. Smith, J. Carroll, M. A. Febbraio, M. Shimada, D. L. Russell, R. L. Robker, Female reproductive life span is extended by targeted removal of fibrotic collagen from the mouse ovary. Sci Adv 8, eabn4564 (2022).

15. Z. Lin, Y. Li, Y. Zhao, D. Liu, S. Deng, J. Gu, Y. Li, X. Zhao, P. Wu, Y. Xiao, J. Su, Y. Sun, Y. Zhang, Y. L. Lee, Y. Sato, H. Zeng, H. Lu, J. Zhang, J. K. Y. Ko, J. Zhao, K. Kawamura, E. H. Y. Ng, S. Jiang, Y. Li, X. Xia, K. K. L. Chan, W. S. B. Yeung, T. R. Wang, K. Liu, Antifibrotic drug finerenone restores fertility in premature ovarian insufficiency. Science 391, eadz4075 (2026).

16. E. R. West, M. Xu, T. K. Woodruff, L. D. Shea, Physical properties of alginate hydrogels and their effects on in vitro follicle development. Biomaterials 28, 4439–48 (2007).

17. S. Pietroforte, F. Amargant, Increased stiffness mimicking ovarian aging induces a fibroinflammatory response in follicles and impairs oocyte quality. Reproduction 171 (2026).

18. R. O. Hynes, A. Naba, Overview of the Matrisome—An Inventory of Extracellular Matrix Constituents and Functions. Cold Spring Harb. Perspect. Biol. 4, a004903 (2012).

19. A. Naba, K. R. Clauser, R. O. Hynes, Enrichment of Extracellular Matrix Proteins from Tissues and Digestion into Peptides for Mass Spectrometry Analysis. J Vis Exp Jove, e53057 (2015).

20. C. R. Below, J. Kelly, A. Brown, J. D. Humphries, C. Hutton, J. Xu, B. Y. Lee, C. Cintas, X. Zhang, V. Hernandez-Gordillo, L. Stockdale, M. A. Goldsworthy, J. Geraghty, L. Foster, D. A. O’Reilly, B. Schedding, J. Askari, J. Burns, N. Hodson, D. L. Smith, C. Lally, G. Ashton, D. Knight, A. Mironov, A. Banyard, J. A. Eble, J. P. Morton, M. J. Humphries, L. G. Griffith, C. Jørgensen, A microenvironment-inspired synthetic three-dimensional model for pancreatic ductal adenocarcinoma organoids. Nat Mater, 1–10 (2021).

21. X. Shao, C. D. Gomez, N. Kapoor, J. M. Considine, C. Grams, Y. (Tom) Gao, A. Naba, MatrisomeDB 2.0: 2023 updates to the ECM-protein knowledge database. Nucleic Acids Res. 51, D1519–D1530 (2022).

22. X. Shao, I. N. Taha, K. R. Clauser, Y. T. Gao, A. Naba, MatrisomeDB: the ECM-protein knowledge database. Nucleic Acids Res 48, D1136–D1144 (2020).

23. H. M. Kinnear, C. E. Tomaszewski, F. L. Chang, M. B. Moravek, M. Xu, V. Padmanabhan, A. Shikanov, The ovarian stroma as a new frontier. Reproduction 160, R25– R39 (2020).

24. C. B. Berkholtz, B. E. Lai, T. K. Woodruff, L. D. Shea, Distribution of extracellular matrix proteins type I collagen, type IV collagen, fibronectin, and laminin in mouse folliculogenesis. Histochem Cell Biol 126, 583–92 (2006).

25. K. Dzobo, C. Dandara, The Extracellular Matrix: Its Composition, Function, Remodeling, and Role in Tumorigenesis. Biomimetics 8, 146 (2023).

26. L. Wullkopf, A.-K. V. West, N. Leijnse, T. R. Cox, C. D. Madsen, L. B. Oddershede, J. T. Erler, Cancer cells’ ability to mechanically adjust to extracellular matrix stiffness correlates with their invasive potential. Mol. Biol. Cell 29, 2378–2385 (2018).

27. A. Jaeschke, M. S. Hepburn, A. Mowla, B. F. Kennedy, C. J. Chan, Three-dimensional quantitative micro-elastography reveals alterations in spatial elasticity patterns in murine ovaries during ageing. *Commun*. Biol. 8, 1409 (2025).

28. S. J. A. Aper, A. C. C. van Spreeuwel, M. C. van Turnhout, A. J. van der Linden, P. A. Pieters, N. L. L. van der Zon, S. L. de la Rambelje, C. V. C. Bouten, M. Merkx, Colorful Protein-Based Fluorescent Probes for Collagen Imaging. PLoS ONE 9, e114983 (2014).

29. T. I. R. Hopkins, V. L. Bemmer, S. Franks, C. Dunlop, K. Hardy, I. E. Dunlop, Micromechanical mapping of the intact ovary interior reveals contrasting mechanical roles for follicles and stroma. Biomaterials 277, 121099 (2021).

30. I. M. Moya, G. Halder, Hippo–YAP/TAZ signalling in organ regeneration and regenerative medicine. Nat Rev Mol Cell Bio 20, 211–226 (2019).

31. S. Pietroforte, M. Plough, F. Amargant, Age-associated increased stiffness of the ovarian microenvironment impairs follicle development and oocyte quality and rapidly alters follicle gene expression. doi: 10.1101/2024.06.09.598134 (2024).

32. S. Granados-Aparici, Q. Yang, H. Clarke, SMAD4 promotes somatic-germLine contact during murine oocyte growth. doi: 10.7554/elife.91798.2 (2024).

33. J. Dong, D. F. Albertini, K. Nishimori, T. R. Kumar, N. Lu, M. M. Matzuk, Growth differentiation factor-9 is required during early ovarian folliculogenesis. Nature 383, 531–535 (1996).

34. A. Abuammah, N. Maimari, L. Towhidi, J. Frueh, K. Y. Chooi, C. Warboys, R. Krams, New developments in mechanotransduction: Cross talk of the Wnt, TGF-β and Notch signalling pathways in reaction to shear stress. Curr. Opin. Biomed. Eng. 5, 96–104 (2018).

35. C. P. Neu, A. Khalafi, K. Komvopoulos, T. M. Schmid, A. H. Reddi, Mechanotransduction of bovine articular cartilage superficial zone protein by transforming growth factor β signaling. Arthritis Rheum. 56, 3706–3714 (2007).

36. X. Tian, A. N. Halfhill, F. J. Diaz, Localization of phosphorylated SMAD proteins in granulosa cells, oocytes and oviduct of female mice. Gene Expr. Patterns 10, 105–112 (2010).

37. A. H. Lin, J. Luo, L. H. Mondshein, P. ten Dijke, D. Vivien, C. H. Contag, T. Wyss- Coray, Global Analysis of Smad2/3-Dependent TGF-β Signaling in Living Mice Reveals Prominent Tissue-Specific Responses to Injury. J. Immunol. 175, 547–554 (2005).

38. M. Giovanni, N. M. F., A. Sandro, D. S. Antonio, F. M. C., C. Fabiana, S. M. L., A. Alessandro, C. Flavio, S. G. C., R. Francesca, V. Maurizio, A. Raja, B. Fabrizio, O. Sara, F. Maria, C. G. R., B. Livia, S. Vincenzo, P. Roberta, O. Ambrogio, P. Francesco, Mongersen, an Oral SMAD7 Antisense Oligonucleotide, and Crohn’s Disease. N. Engl. J. Med. 372, 1104–1113 (2015).

39. S. Ardizzone, G. Bevivino, G. Monteleone, Mongersen, an oral Smad7 antisense oligonucleotide, in patients with active Crohn’s disease. Ther. Adv. Gastroenterol. 9, 527– 532 (2016).

40. M. Wu, W. Tang, Y. Chen, L. Xue, J. Dai, Y. Li, X. Zhu, C. Wu, J. Xiong, J. Zhang, T. Wu, S. Zhou, D. Chen, C. Sun, J. Yu, H. Li, Y. Guo, Y. Huang, Q. Zhu, S. Wei, Z. Zhou, M. Wu, Y. Li, T. Xiang, H. Qiao, S. Wang, Spatiotemporal transcriptomic changes of human ovarian aging and the regulatory role of FOXP1. *Nat*. Aging 4, 527–545 (2024).

41. G. Nagamatsu, S. Shimamoto, N. Hamazaki, Y. Nishimura, K. Hayashi, Mechanical stress accompanied with nuclear rotation is involved in the dormant state of mouse oocytes. Sci Adv 5, eaav9960 (2019).

42. J. E. Hornick, F. E. Duncan, L. D. Shea, T. K. Woodruff, Isolated primate primordial follicles require a rigid physical environment to survive and grow in vitro. Hum Reprod 27, 1801–10 (2012).

43. X. Zhao, S. Zhang, S. Gao, H.-M. Chang, P. C. K. Leung, J. Tan, A Novel Three- Dimensional Follicle Culture System Decreases Oxidative Stress and Promotes the Prolonged Culture of Human Granulosa Cells. ACS Appl. Mater. Interfaces 15, 15084–15095 (2023).

44. P. K. Kreeger, J. W. Deck, T. K. Woodruff, L. D. Shea, The in vitro regulation of ovarian follicle development using alginate-extracellular matrix gels. Biomaterials 27, 714–23 (2006).

45. A. R. Sun, M. F. H. RamLi, X. Shen, K. K. Ramakanth, D. Chen, Y. Hu, P. Vidyasekar, R. S. Foo, Y. Long, J. Zhu, M. Ackers-Johnson, J. L. Young, Hybrid hydrogel–extracellular matrix scaffolds identify biochemical and mechanical signatures of cardiac ageing. Nat. Mater. 24, 1489–1501 (2025).

46. S. Granados-Aparici, A. Volodarsky-Perel, Q. Yang, S. Anam, T. Tulandi, W. Buckett, W.-Y. Son, G. Younes, J.-T. Chung, S. Jin, M.-E. Terret, H. J. Clarke, MYO10 promotes transzonal projection-dependent germ line-somatic contact during mammalian folliculogenesis. Biol. Reprod. 107, 474–487 (2022).

47. Y. Ishikawa-Yamauchi, C. Emori, H. Mori, T. Endo, K. Kobayashi, Y. Watanabe, H. Sagara, T. Nagata, D. Motooka, A. Ninomiya, M. Ozawa, M. Ikawa, Age-associated aberrations of the cumulus-oocyte interaction and in the zona pellucida structure reduce fertility in female mice. *Commun*. Biol. 7, 1692 (2024).

48. C. A. Doherty, F. Amargant, S. Y. Shvartsman, F. E. Duncan, E. R. Gavis, Bidirectional communication in oogenesis: a dynamic conversation in mice and Drosophila. Trends Cell Biol, doi: 10.1016/j.tcb.2021.11.005 (2021).

49. S. Granados-Aparici, Q. Yang, H. Clarke, SMAD4 promotes somatic-germLine contact during oocyte growth. doi: 10.7554/elife.91798.1 (2024).

50. M. Chen, C. He, K. Zhu, Z. Chen, Z. Meng, X. Jiang, J. Cai, C. Yang, Z. Zuo, Resveratrol ameliorates polycystic ovary syndrome via transzonal projections within oocyte-granulosa cell communication. Theranostics 12, 782–795 (2022).

51. D. Cheng, Y.-H. Chen, X.-P. Jiang, L.-Y. Li, Y. Tan, M. Li, Z.-C. Mo, Transzonal Projections and Follicular Development Abnormalities in Polycystic Ovary Syndrome. Prog. Biochem. Biophys. 52, 2499–2511 (2025).

52. H. E. Sumbul, B. S. Avci, M. Bankir, B. C. Pekoz, E. Gulumsek, A. S. Koc, Ovarian Stiffness Is Significantly Increased in Polycystic Ovary Syndrome and Related With Anti- Mullerian Hormone. Ultrasound Q. 38, 83–88 (2022).

53. Z. Qin, W. Xia, G. J. Fisher, J. J. Voorhees, T. Quan, YAP/TAZ regulates TGF-β/Smad3 signaling by induction of Smad7 via AP-1 in human skin dermal fibroblasts. Cell Commun. Signal. 16, 18 (2018).

54. K. W. Yong, Y. Li, F. Liu, B. Gao, T. J. Lu, W. A. B. W. Abas, W. K. Z. W. Safwani, B. Pingguan-Murphy, Y. Ma, F. Xu, G. Huang, Paracrine Effects of Adipose-Derived Stem Cells on Matrix Stiffness-Induced Cardiac Myofibroblast Differentiation via Angiotensin II Type 1 Receptor and Smad7. Sci. Rep. 6, 33067 (2016).

55. M. T. Zenzes, R. Bielecki, Nicotine-induced Disturbances of Meiotic Maturation in Cultured Mouse Oocytes: Alterations of Spindle Integrity and Chromosome Alignment. Tob. Induc. Dis. 2, 151 (2004).

56. Z. Feng, P. Fang, H. Zheng, X. Zhang, DEP2: an upgraded comprehensive analysis toolkit for quantitative proteomics data. Bioinformatics 39, btad526 (2023).

57. A. Naba, K. R. Clauser, H. Ding, C. A. Whittaker, S. A. Carr, R. O. Hynes, The extracellular matrix: Tools and insights for the “omics” era. Matrix Biol. 49, 10–24 (2016).

58. C. Chen, H. Chen, Y. Zhang, H. R. Thomas, M. H. Frank, Y. He, R. Xia, TBtools: An Integrative Toolkit Developed for Interactive Analyses of Big Biological Data. Mol Plant 13, 1194–1202 (2020).

